# Live-cell PAINT microscopy of protein targets *via* fluorogenic exchangeable HaloTag ligands

**DOI:** 10.64898/2026.07.16.738981

**Authors:** Tong Zhan, Nels C. Gerstner, Julia G. Martin, Jack T. McCann, Marissa C. Lynch, Hsin-Jung Wu, Ke Xu, Evan W. Miller

**Affiliations:** Departments of Chemistry, University of California, Berkeley 94720-1460; Departments of Molecular & Cell Biology, University of California, Berkeley 94720-1460; Departments of Helen Wills Neuroscience Institute, University of California, Berkeley 94720-1460

## Abstract

We report a generalizable method for single-molecule localization super-resolution microscopy in living cells. Live-cell PAINT (points accumulation for imaging in nanoscale topography) and single-molecule diffusivity mapping (SM*d*M) of intracellular protein targets are achieved by pairing exceptionally fluorogenic bis-trifluoromethyl rhodamine (BF) dyes with reversible HaloTag ligands and widely available HaloTag fusion proteins. This far-red small-molecule update to protein-based PAINT is readily incorporated into existing super-resolution microscopy workflows: wash-free staining and imaging are achieved for cell lines and primary neuron cultures for diverse protein targets. Pairing with photoconvertible fluorescent proteins further enables simultaneous two-color live-cell super-resolution microscopy and SM*d*M.

---

Single-molecule localization microscopy (SMLM) offers a powerful yet highly accessible approach to super-resolution microscopy (SRM).^1-4^ In SMLM, the super-localized positions of millions of single molecules are mass-accumulated in the wide field over thousands of camera frames to construct images at ∼10 nm spatial resolution. Extending single-molecule measurements to additional dimensions further enables multidimensional SRM,^5^ *e*.*g*., single-molecule displacement/diffusivity mapping (SM*d*M)^6-7^ for super-resolution diffusion mapping.

The performance of SMLM depends critically on fluorophores that both provide strong single-molecule fluorescence over the background and exhibit long-lasting stochastic switching between an emitting (fluorescent) state and a non-emitting (dark) state.^8-9^ Among the different approaches to achieve these goals,^4, 8-9^ PAINT, or points accumulation for imaging nanoscale topography,^10^ stands out because of its compatibility with a range of fluorophores, independence from buffer additives, and low illumination intensity requirements. PAINT sustains long-lasting single-molecule fluorescence switching by the reversible binding of fluorophores partitioning from a pool of latent, dark states to target structures. This was originally accomplished with environmentally sensitive dyes partitioning into lipid membranes,^10^ and then extended to fluorophores binding to cell-surface proteins^11^ and nucleic acids.^12^ For intracellular protein targets, DNA-PAINT provides a versatile solution in fixed cells by utilizing complementary DNA strands tagged to antibodies or nanobodies.^13-14^ PAINT of protein targets in living cells remains challenging, typically relying on fluorescent protein-peptide interactions^15-18^ or fluorogenic binding of fluorescent-protein derivatives.^19-21^ Neither approach uses modern synthetic rhodamine fluorophores, which are bright and available in long wavelengths to facilitate multicolor imaging and reduce background.^22-23^

Recent, elegant approaches^24-25^ use the self-labeling protein, HaloTag, which forms a covalent bond with a chloroalkane ligand, to access bright rhodamine dyes for PAINT. To achieve reversibility, two approaches can be used. First, HaloTag itself can be re-engineered to hydrolyze the enzyme-substrate intermediate.^24^ This allows the use of typical chloroalkane HaloTag ligands (HTL) with a re-engineered enzyme. The second approach is to use the original HaloTag, but pair it with a pseudo-halide ligand which cannot form a covalent bond with the HaloTag protein, creating a reversible, exchangeable HaloTag ligand (xHTL).^25^ While the xHTL approach enables PAINT in fixed cells, live-cell PAINT remains elusive.^25-26^ Dyes with high fluorogenicity indices appeared to perform better in fixed cell PAINT. Therefore, we hypothesized that reducing the background fluorescence from unbound dye might improve the performance of HaloTag-based PAINT (**Scheme 1a**).

We recently discovered that replacing the classic bridging oxygen atom at the 10’ position of rhodamine dyes with a bis-trifluoromethyl group results in a ∼90 nm red-shift into the farred region of the spectrum. These new **<u>b</u>**is-**f**luorinated dyes (**<u>B</u>**erkeley**<u>F</u>**luor, or BF dyes) are unique because the close-open equilibrium (K_LZ_) is very low (<2.2 × 10^−3^),^27^ strongly favoring the non-absorbent, non-fluorescent closed form (**Scheme 1a,b**). We previously showed^27^ that when BF dyes are paired with the chloroalkane HaloTag ligand (HTL), BF dyes become highly fluorogenic upon binding HaloTag proteins, resulting in a 25-50× in *in vitro* fluorescence and a 10,000× increase in cellular fluorescence, a ∼20× improvement in their fluorogenicity compared to their analogous silicon rhodamine (SiR) counterparts.^27^ The improved fluorogenicity of BF dyes comes from a substantially lower amount of the open, zwitterionic (Z), form of the dye in the absence of HaloTag (**Scheme 1b**).^27^ Owing to the high fluorogenicity of irreversible chloroalkane-based BF-HTLs, we reasoned that BF dyes could be converted into probes for PAINT microscopy, by taking advantage of exchangeable HaloTag ligands (xHTLs). BF dyes coupled to an xHTL, like **BF-646-xHTL(T5)** (**Scheme 1b**), could reversibly bind HaloTag fusion proteins to enable PAINT microscopy.

**Scheme 1.**
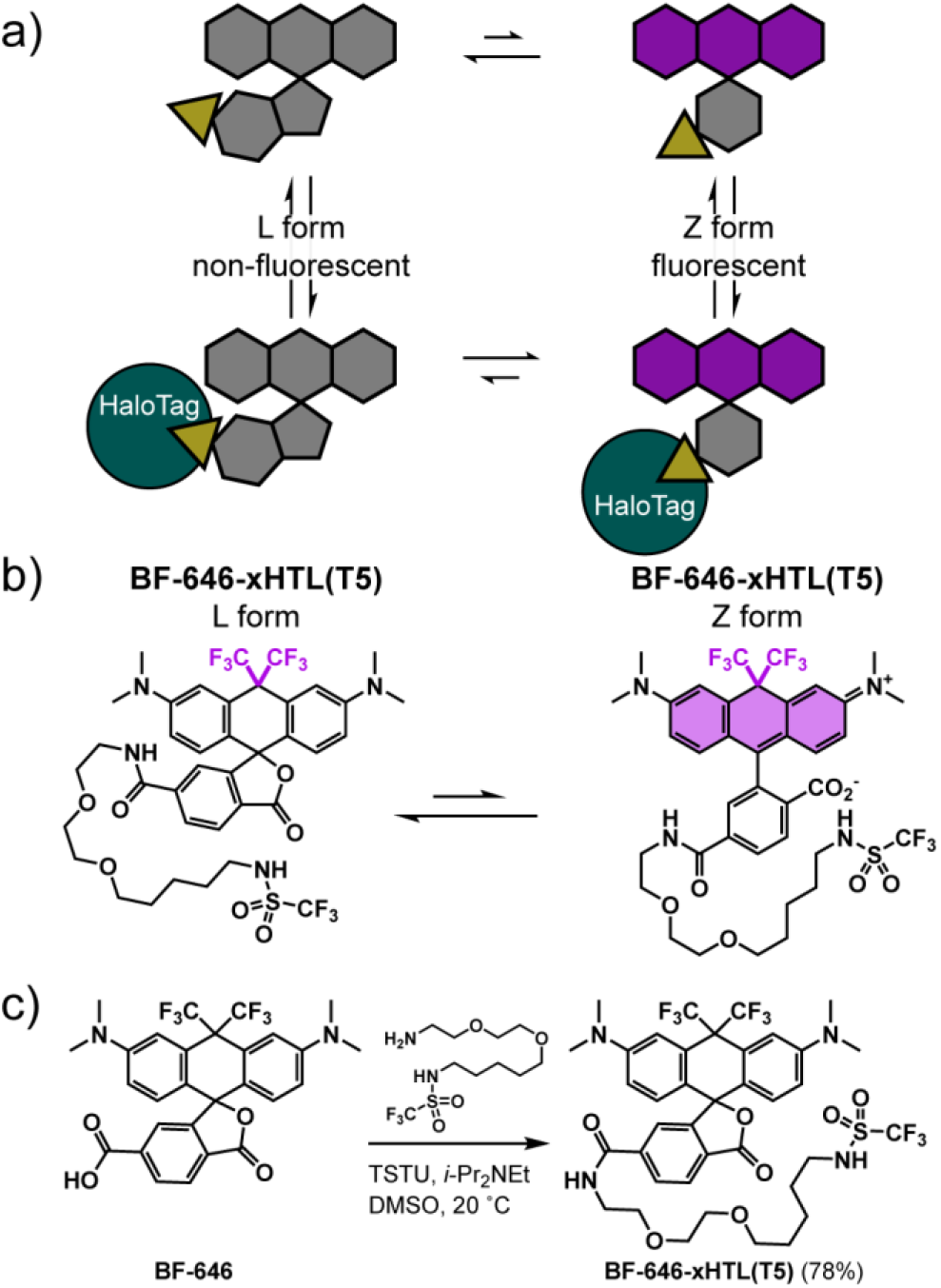
BerkeleyFluor exchangeable HaloTag ligands. **a)** In the absence of HaloTag, BF-646 favors the closed, lactone form over the open, colored, and fluorescent zwitterionic form. Reversible binding to HaloTag (aqua) shifts the equilibrium to favor the open form. **b)** Structures of **BF-646-xHTL(T5)** in the lactone (L) and zwitterionic (Z) forms. **c)** Synthesis of **BF-646-xHTL(T5)**.

**BF-646-xHTL(T5)** uses the previously reported triflimide-based xHTL ligand, “T5”,^25^ and can be synthesized in a single step via a TSTU-mediated coupling between the previously reported BF-646-acid^27^ and T5 xHTL ligand^25^ in 78% yield (**Scheme 1c, S1**). **BF-646-xHTL(T5)** is a ligand for HaloTag and shows strong fluorogenicity upon binding to purified HaloTag protein (**Figure 1a**). When bound to HaloTag, **BF-646-xHTL(T5)** has an absorbance maximum of 652 nm, an emission maximum at 665 nm, and a fluorescence quantum yield (Φ_fl_) of 27% (**Figure 1c**). The optical properties of **BF-646-xHTL(T5)** in the presence of HaloTag are nearly identical to the previously reported BF-646-HTL (irreversible chloroalkane ligand): 651 nm absorbance, 664 nm emission, and a 35% fluorescence quantum yield.^27^ The absorbance of **BF-646-xHTL(T5)** increases 14× in the presence of HaloTag; the emission by a factor of 15× (**Figure 1c**),

**Figure 1.**
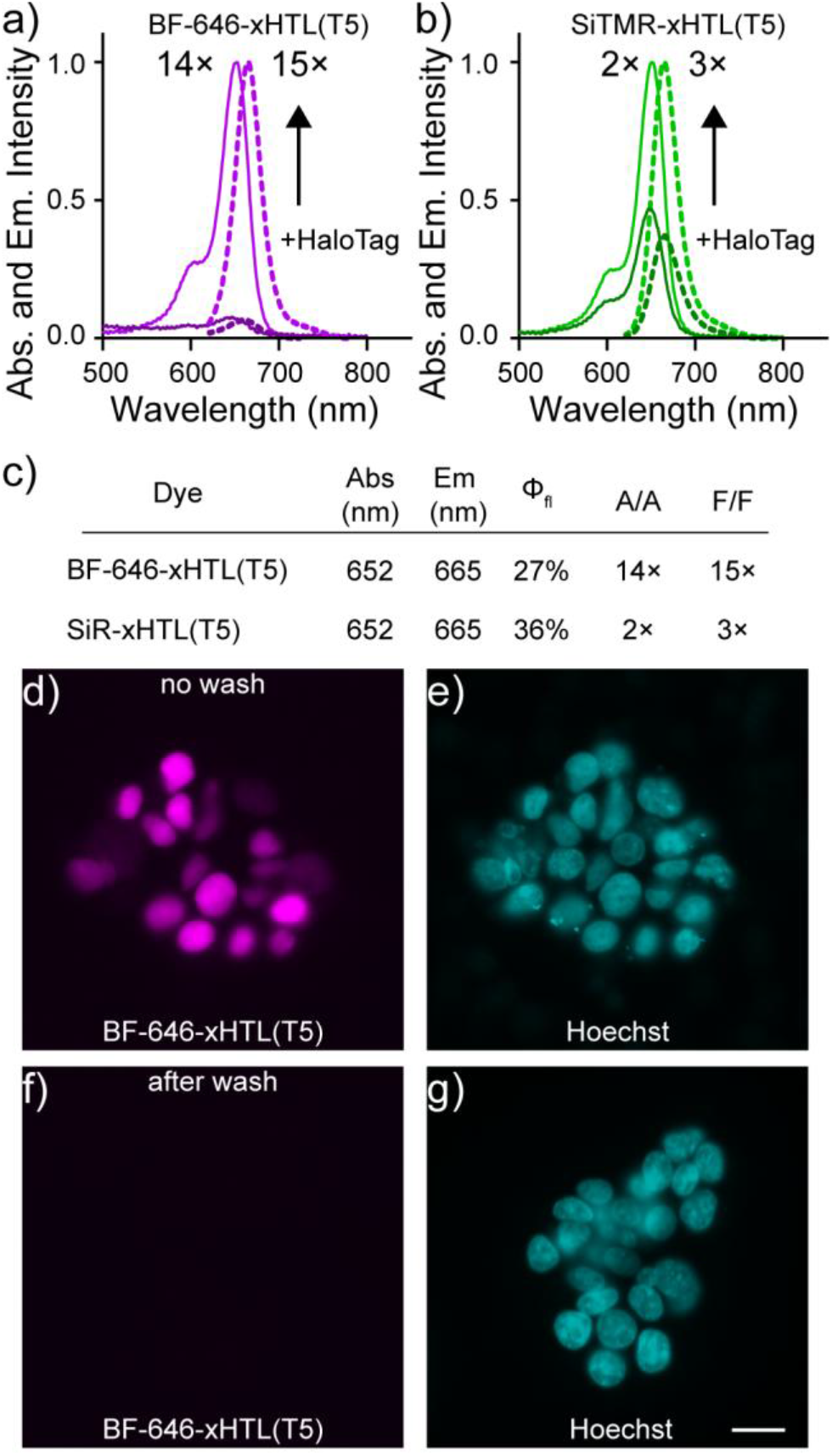
*In vitro* and cellular characterization of BF-646-xHTL(T5). Plots of normalized absorbance (solid line) and emission intensity (dashed line) vs wavelength for **a)** BF-646-xHTL(T5) and **b)** the corresponding SiR-xHTL(T5) in the absence or presence of purified HaloTag protein (1 μM dye, 2 μM HaloTag). **c)** Summarized optical properties of compounds. Optical properties were measured in Dulbecco’s modification to phosphate buffered saline, pH 7.2 (dPBS). A/A and F/F are the ratio of peak absorbance (A) or fluorescence emission (F) with and without HaloTag. Φfl is the fluorescence quantum yield. Epifluorescence images of HEK293T cells expressing nuclear-localized HaloTag and stained with **d)** BF-646-xHTL(T5) (200 nM) and co-stained with **e)** nuclear dye Hoechst 33342. Epifluorescence images of HEK293T cells expressing nuclear-localized HaloTag and stained with **f)** BF-646-xHTL(T5) (200 nM) followed by washing in dye-free buffer and co-stained with **g)** nuclear dye Hoechst 33342.

which is 5 to 7× better than the corresponding, previously reported, SiR-xHTL(T5) (2× for absorbance and 3× for emission, **Figure 1b**).^25^

**BF-646-xHTL(T5)** binds HaloTag in living cells. HEK293T cells expressing nuclear-localized HaloTag show bright, far-red fluorescence localized to the nucleus when incubated in the presence of 200 nM **BF-646-xHTL(T5)** (**Figure 1d-e**). Cellular binding of HaloTag by **BF-646-xHTL(T5)** is reversible: when **BF-646-xHTL(T5)** is removed from the buffer, the fluorescence intensity drops by over 200× (**Figure 1f-g** and **Figure S1**). The high brightness, large HaloTag-dependent fluorogenicity, and reversible binding of cellular HaloTag make BF-xHTLs good candidates for HaloTag-PAINT in living cells.

**BF-646-xHTL(T5)** enables PAINT of protein targets in live cells. When imaged with a typical SMLM setup at 56 frames per second (fps), keeping 2.5 nM **BF-646-xHTL(T5)** in the medium of living COS-7 cells expressing HaloTag-Sec61β that targets the endoplasmic reticulum (ER) membrane produces excellent single-molecule images with high brightness and minimal background (**Figure 2a**), attributable to the high fluorogenicity of **BF-646-xHTL(T5)** upon binding HaloTag. The single-molecule fluorescence of **BF-646-xHTL(T5)** exhibits robust on-off switching through HaloTag binding/unbinding, maintaining a good single-molecule density of ∼4 µm^-2^s^-1^ with no discernible drops over 50,000 frames (15 min) (**Figure 2b** and **Movie S1**). In comparison, under identical conditions, SiR-xHTL(T5) exhibits a rapid drop in single-molecule counts from ∼4 µm^-2^s^-1^ to ∼0.7 µm^-2^s^-1^ within the first ∼1,000 frames (**Figure 2b** and **Movie S1**). This result may be explained by the lower fluorogenicity and higher K_LZ_ of SiR-xHTL(T5): the unbound dye is excited and photobleached before it can diffuse to the HaloTag. ∼2,000 photons were detected for each **BF-646-xHTL(T5)** molecule per frame, substantially higher than ∼1,200 for SiR-xHTL(T5) (**Figure 2c**). This high brightness implies localization precisions of ∼10 nm in standard deviation.

**Figure 2.**
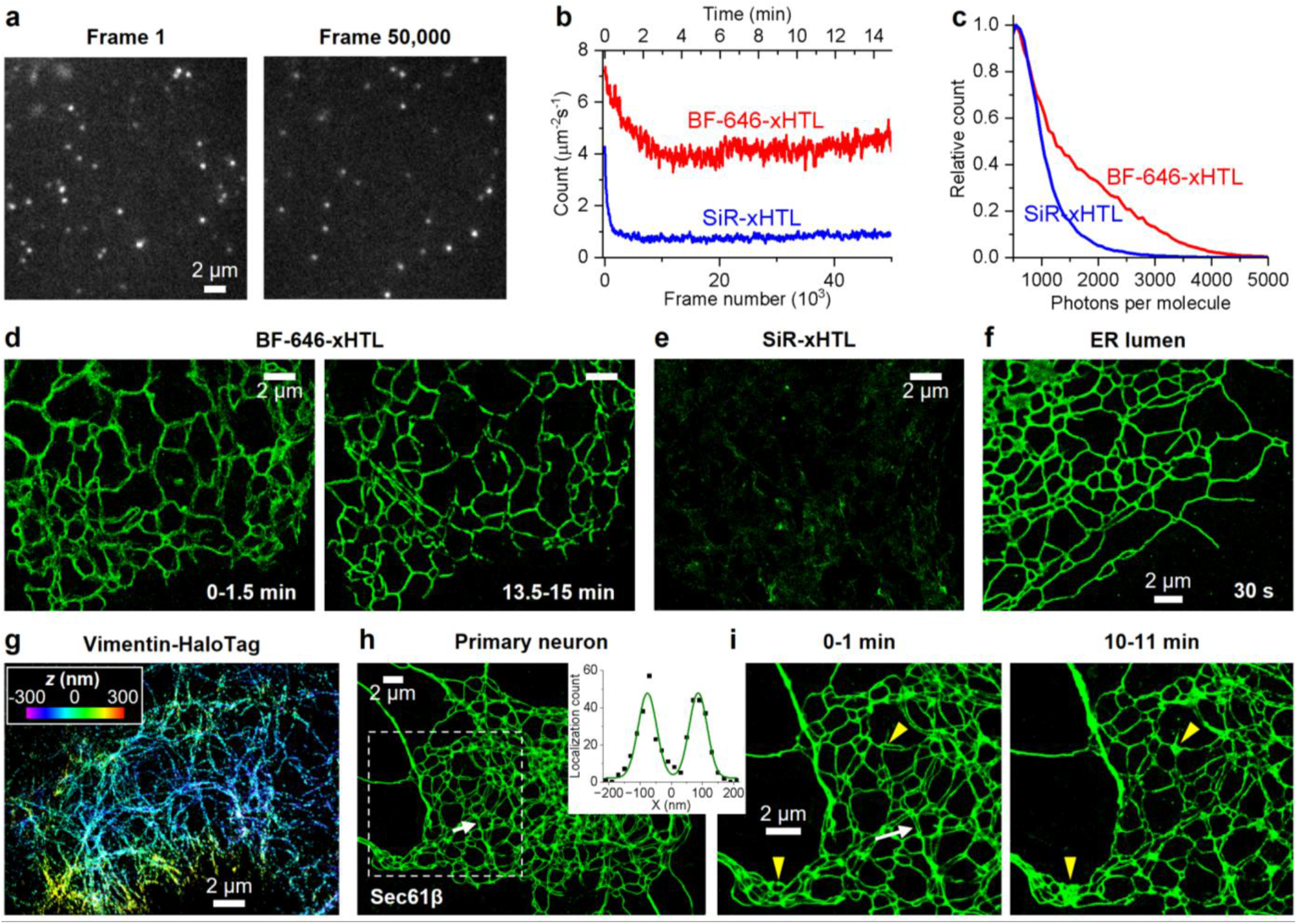
BF-646-xHTL(T5) enables PAINT super-resolution microscopy of protein targets in living cells. (**a**) Representative single-molecule images recorded when 2.5 nM BF-646-xHTL(T5) is kept in the medium of living COS-7 cells expressing HaloTag-Sec61β targeting the endo-plasmic reticulum (ER) membrane. Left and right panels are from the beginning and after continuous recording 50,000 frames at 56 frames per second (fps), respectively. (**b**) Time-dependent counts of the detected BF-646-xHTL(T5) single molecules per μm^2^ per second over the recorded 50,000 frames (red), compared to SiR-xHTL(T5) under the same conditions in another sample (blue). (**c**) Distributions of photon counts for single molecules recorded in each frame for the two dyes. (**d**) Live-cell PAINT images generated from BF-646-xHTL(T5) molecules detected in the initial (left) and final (right) 5,000 frames, corresponding to 0-1.5 min and 13.5-15 min, respectively. (**e**) Live-cell PAINT image generated from SiR-xHTL(T5) molecules detected in the initial 5,000 frames. (**f**) Live-cell PAINT image with BF-646-xHTL(T5) labeling ER-HaloTag diffusing in the ER lumen, from single molecules accumulated over 30 s at 109 fps. (**g**) 3D-PAINT image with BF-646-xHTL(T5) labeling vimentin-HaloTag in a fixed COS-7 cell. Color presents depth (z). (**h**) Live-cell PAINT image of BF-646-xHTL(T5) labeling HaloTag-Sec61β in a cultured primary hippocampal neuron. Inset: intensity profile (as count of single-molecule localizations) at the white arrows in (**h**,**i**), resolving 68 nm ER-tubule widths and a 160 nm center-to-center distance between two adjacent ER tubules. (**i**) Zoom-ins of the box in (**h**), reconstructed from 0-1 min (Left) and 10-11 min (Right) of the data. Yellow arrows indicate regions of tube-to-sheet ER transitions.

The robust single-molecule blinking and high brightness of **BF-646-xHTL(T5)** allowed us to construct time series of live-cell PAINT super-resolution images, *e*.*g*., resolving the intricate and dynamic ER networks at a 1.5-min temporal resolution, with no degradation in image quality over time (**Figure 2d, Figure S2**, and **Movie S2**). The high fluorogenicity of **BF-646-xHTL(T5)** further yielded excellent target specificity under the no-wash condition, showing minimal signal outside the ER network. In contrast, SiR-xHTL(T5) did not resolve the ER structure under identical conditions (**Figure 2e**) due to the low count of detected molecules.

**BF-646-xHTL(T5)** is compatible with labeling a variety of cellular structures. **BF-646-xHTL(T5)** labels soluble ER-HaloTag in the lumen of the ER in COS-7 cells, for which 109-fps recording under 1-ms stroboscopic excitation allowed PAINT image series to be constructed at a 30-s temporal resolution (**Figure 2f, Figure S3**, and **Movie S3**). We further demonstrated the application of **BF-646-xHTL(T5)** to PAINT imaging of vimentin-HaloTag in both live (**Figure S4**) and fixed cells, including 3D-PAINT (**Figure 2g**).

The high brightness and fluorogenicity of **BF-646-xHTL(T5)** enable PAINT in live hippocampal neurons. Live-cell PAINT imaging in rat hippocampal neurons expressing ER-membrane-associated HaloTag-Sec61β reveals a fine ER network that dynamically remodels on the 1-min time scale (**Figure 2h,i, Figure S5**, and **Movie S4**). Intensity profiles across two adjacent ER tubules resolved a 68 nm tubule width and a 160 nm center-to-center distance (**Figure 2h inset**, white arrows), substantially below the ∼250 nm diffraction limit. This high resolution helped resolve dynamic structural changes like tubule-to-sheet transitions (yellow arrowheads in **Figure 2i**).

The far-red emission of **BF-646-xHTL(T5)** further provides an exceptional opportunity for multiplexing. Modern live-cell SMLM often relies on bright photoconvertible fluorescent proteins such as Eos and Dendra to achieve photoactivated localization microscopy (PALM)^4, 8^-^9, 28^, but their crossover between photoconverting and imaging light in both the green and red channels precludes their concurrent usage. The far-red excitation and emission profile of **BF-646-xHTL(T5)** allows simultaneous use with Eos or Dendra fluorescent proteins for two-color, live-cell PALM-PAINT.

With mEos4b-TOM20 and HaloTag-Sec61β co-expressed in a COS-7 cell, **BF-646-xHTL(T5)** enables concurrent two-color PALM-PAINT imaging of mitochondria and ER in the living cell (**Figure 3a**,**b**) through alternating 561 nm and 647 nm excitations between frames. Nanoscale dynamic interactions between the two targets were captured at a 1 min temporal resolution (**Figure 3b**,**c**) with no signal crosstalk (**Figure 3d**), allowing observation of mitochondrial fission preceded by encircling by ER (Arrowheads in **Figure 3b**,**d**). **BF-646-xHTL(T5)** maintained a good single-molecule count through HaloTag binding/unbinding under the alternating excitation scheme, which slowly dropped from ∼8 µm^-2^s^-1^ to ∼4 µm^-2^s^-1^ over 140,000 frames (21 minutes) (**Figure 3e**). In comparison, mEos4b showed a sharp drop in single-molecule count in the initial ∼2,000 frames, but the application of a weak 405-nm laser helped sustain a workable single-molecule density (**Figure 3e**) by photoconverting more mEos4b to the 561-nm-excitable state. We thus obtained good PAINT-PALM image series for both targets over >20 min (**Figure 3f, g, Figure S6**, and **Movie S5**). The probe-exchange-based spontaneous maintenance of single-molecule counts for the **BF-646-xHTL(T5)**-labeled target was critical for enabling concurrent two-color SMLM, which allowed the 405-nm laser power to be freely adjusted during the experiment to optimize the mEos4b single-molecule density.

**Figure 3.**
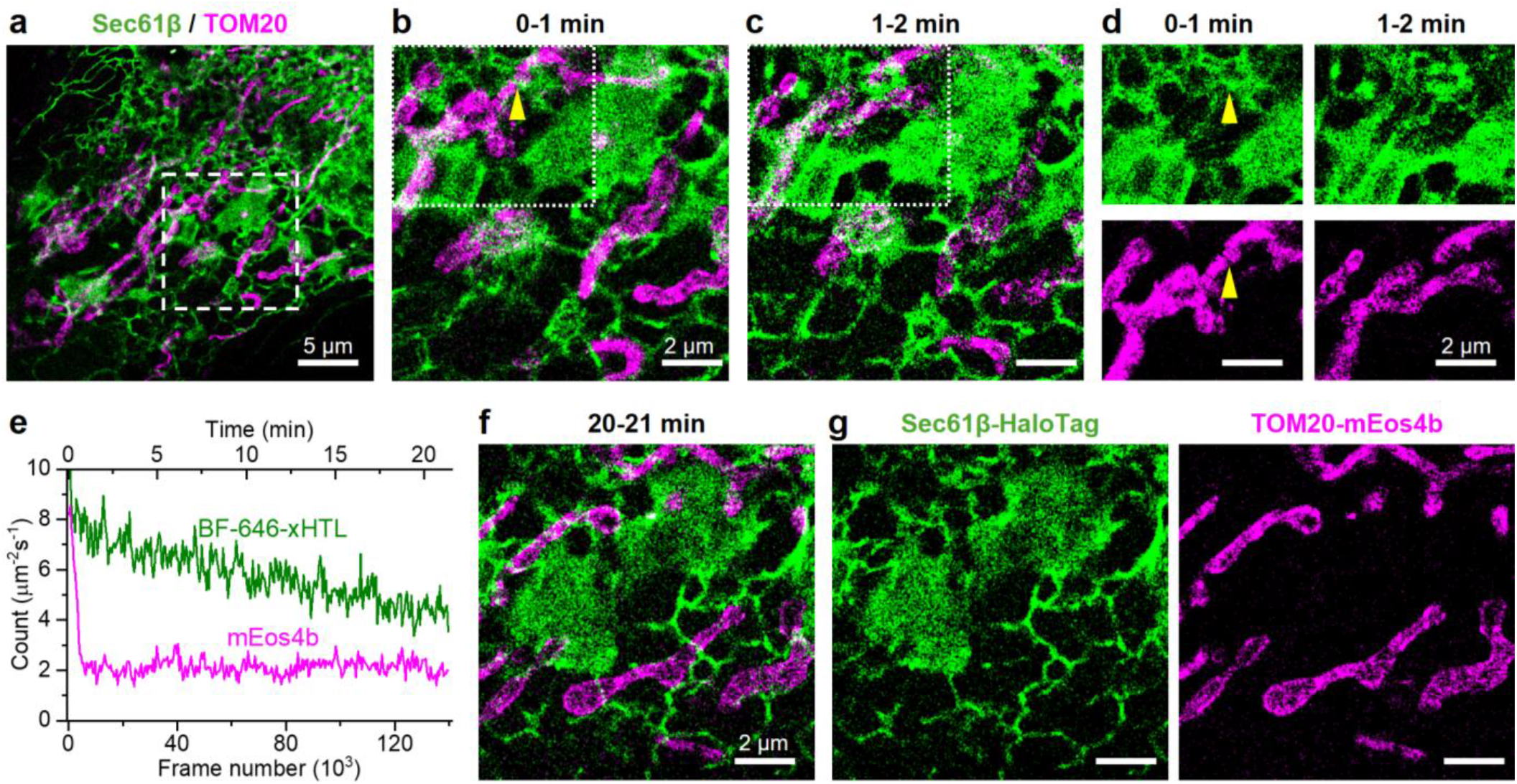
BF-646-xHTL(T5) enables simultaneous two-color PALM-PAINT super-resolution microscopy of protein targets in living cells. (**a**) Concurrent PALM of mEos4b-TOM20 (magenta) and PAINT of BF-646-xHTL(T5)-labeled HaloTag-Sec61β (green) in a living COS-7 cell, achieved by alternating 561 nm and 647 nm excitations between frames when recording at 109 fps. The two-color PALM-PAINT image is reconstructed from 6,540 frames recorded over 1 minute. (**b**) Zoom-in of the box in (a). (**c**) Same as (b) but reconstructed from data recorded in the next minute. (**d**) Separately presented PAINT and PALM images of the two targets for the boxed region in (b,c). (**e**) Time-dependent counts of the detected BF-646-xHTL(T5) (green) and mEos4b (magenta) single molecules per μm^2^ per second for the region shown in (b,c,f,) over the recorded 140,000 frames (21 minutes). (**f**) The same region as (b,c) for data recorded during 20-21 min. (**g**) Separately presented PAINT and PALM images of the two targets in (f).

**BF-646-xHTL(T5)** further enables two-color SM*d*M in living cells. In SM*d*M, closely timed stroboscopic excitation pulses are repeatedly applied across tandem frames to capture transient single-molecule displacements and enable super-resolution diffusion mapping.^6-7^ However, the application of SM*d*M to intracellular protein targets has been limited to fluorescent proteins, which offer low brightness and do not enable two-color experiments.

The SMLM data in Figure 2f on **BF-646-xHTL(T5)**-labeled ER-HaloTag was obtained by repeatedly applying 2-ms-separated excitation pulses across paired frames, enabling SM*d*M analysis. Evaluating the vectorial single-molecule displacements^29^ (**Figure 4a**) for local statistics on a 100 nm spatial grid, SM*d*M shows that **BF-646-xHTL(T5)**-labeled ER-HaloTag diffuses along local ER-tubule orientations (**Figure 4b**) with a relatively uniform diffusion coefficient of ∼8 µm^2^/s (**Figure 4c**), consistent with our previous results^30^ with ER-Dendra2 and other reported diffusion coefficients in the ER lumen (*D* = 5-10 μm^2^/s).^30-32^ The robust single-molecule signal further allowed us to construct time series of SM*d*M diffusion maps at a 2-min resolution (**Figure 4c**,**d, Figure S7** and **Movie S6**).

**Figure 4.**
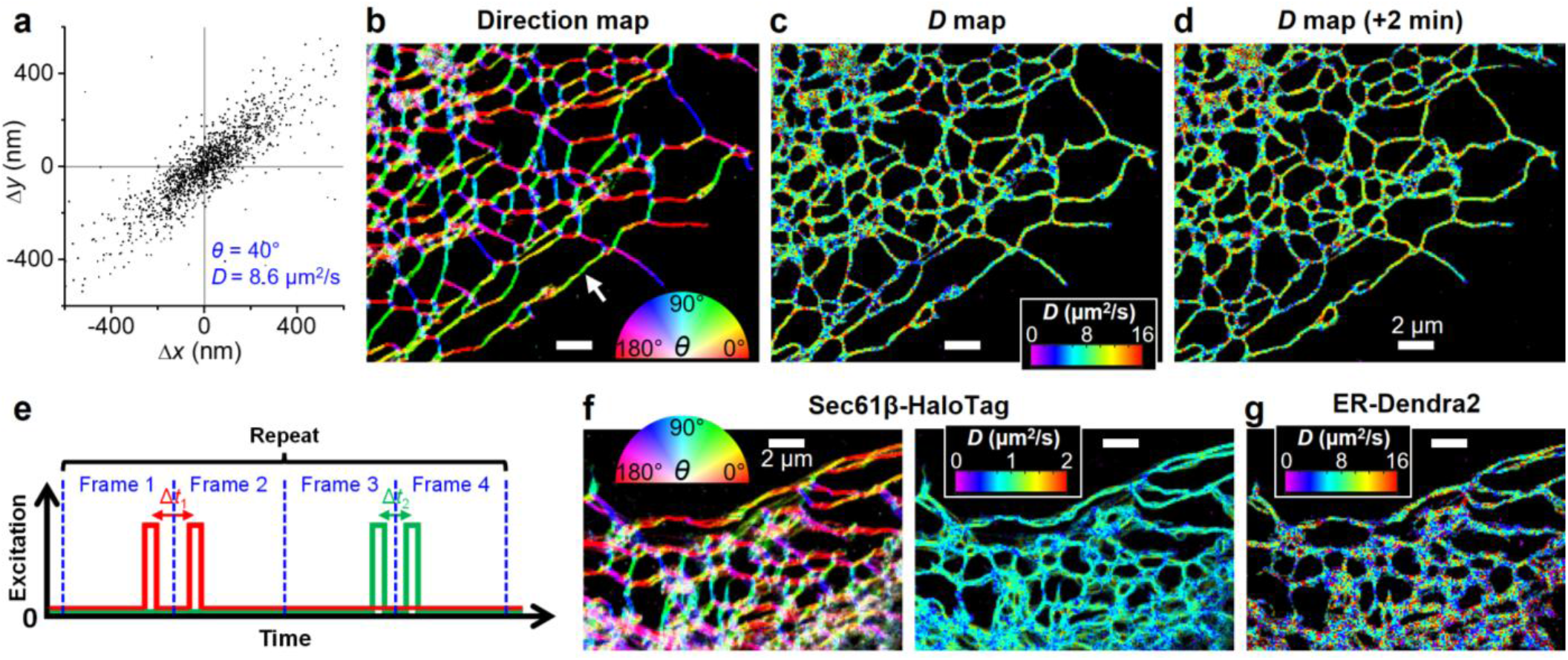
BF-646-xHTL(T5) enables concurrent two-color SM*d*M super-resolution diffusion mapping of protein targets in living cells. (**a**) Plot of SM*d*M-accumulated 2-ms vectorial single-molecule displacements of BF-646-xHTL(T5)-labeled ER-HaloTag diffusing in an ER-tubule segment of a COS-7 cell [white arrows in (b)]. A principal direction *θ* of 40° is assessed, along which the diffusion coefficient is determined to be 8.6 µm^2^/s. (**b**,**c**) Color-coded SM*d*M super-resolution maps of the local principal direction of diffusion (**b**) and diffusion coefficient (**c**) of BF-646-xHTL(T5)-labeled ER-HaloTag in the cell (same cell as Fig. 2f), obtained by binning single-molecule displacements collected over 2 minutes onto a 100-nm spatial grid for local statistics. (**d**) Same as (c) but reconstructed from data recorded in the next 2 minutes. (**e**) Schematics: Concurrent two-color SM*d*M *via* alternating between tandem 647-nm and 561-nm excitation pairs. (**f**,**g**) Concurrent two-color SM*d*M of BF-646-xHTL(T5)-labeled HaloTag-Sec61β in the ER membrane (**f**) and ER-Dendra2 in the ER lumen (**g**) in a living COS-7 cell.

The distinct far-red emission of **BF-646-xHTL(T5)** enabled concurrent two-color SM*d*M when paired with photoconvertible fluorescent proteins. To this end, we devised an excitation scheme in which we alternated between 647-nm and 561-nm tandem pulses across paired frames (**Figure 4e**). By separately analyzing transient single-molecule displacements recorded under the 647-nm and 561-nm excitations, we constructed two-color SM*d*M diffusion maps, resolving contrasting diffusion coefficients between the membrane-localized HaloTag-Sec61β (*D* ∼1 μm^2^/s; **Figure 4f**) and the lumenal ER-Dendra2 (*D* ∼8 μm^2^/s; **Figure 4g**) in the same ER networks.

In conclusion, we have shown that the highly fluorogenic BF dyescan be paired with xHTL ligands and widely available Hal-oTag fusion proteins to enable PAINT and SM*d*M super-resolution microscopy for diverse protein targets in living cells. Wash-free staining and imaging with **BF-646-xHTL(T5)** are achieved for diverse protein targets in immortalized cell lines and sensitive primary neuron cultures. The far-red excitation and emission profiles of BF-646 allow it to be readily incorporated into existing super-resolution workflows for concurrent two-color SMLM and SM*d*M. Together, these characteristics make BF-xHTL a practical method for super-resolution microscopy in any context where HaloTag fusion proteins can be employed.

## Acknowledgements

We acknowledge support from the National Institute of General Medical Sciences of the National Institutes of Health (R35GM153237, EWM; R35GM149349, KX; F32GM139263, NCG; T32GM152983, JTMc, JGM, MCL), and the Agilent Biodesign Program (EWM).

## ASSOCIATED CONTENT

Additional experimental details, including synthetic procedures, additional figures, and spectra..

## ACKNOWLEDGMENT

We acknowledge support from the National Institute of General Medical Sciences of the National Institutes of Health (R35GM153237, EWM; R35GM149349, KX; F32GM139263, NCG; T32GM), and the Agilent Biodesign Program (EWM).

---

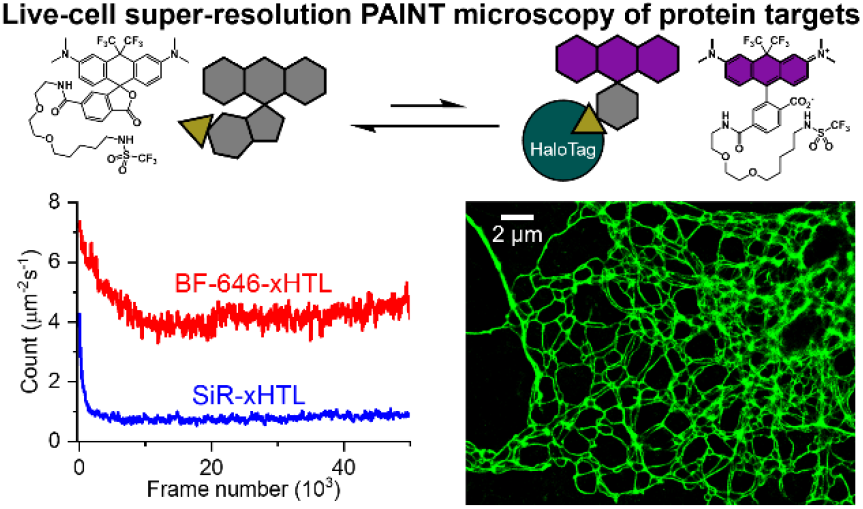

