## Supporting Information for "Live-cell PAINT microscopy of protein targets *via* fluorogenic exchangeable HaloTag ligands"

**Affiliations**:

DOI: https://www.biorxiv.org/content/10.64898/2026.07.16.738981v2

### Chemical Synthesis and Characterization

Chemical reagents and anhydrous solvents in Sureseal bottles were purchased from commercial suppliers and used without further purification. The exceptions were toluene and diethyl ether, which were stored over freshly activated 4Å MS for three days before their use. Thin layer chromatography (TLC) (silica gel, F254, 250 mm) and preparative thin layer chromatography (PTLC) (Silicycle, F254, 1000 mm) were performed on precoated TLC glass plates and were visualized by fluorescence quenching under UV light. Flash column chromatography was performed on Silicycle Silica Flash F60 (230–400 Mesh) for normal phase or 60 RP-18 (200-400 mesh) for reverse phase, using a forced flow of air at 0.5–1.0 bar. NMR spectra were recorded on BrukerAV-300 MHz, BrukerAVB-400 MHz, Bruker AVQ-400 MHz, Bruker NEO-500 MHz, and Bruker AV-600 NMR spectrometers. Chemical shifts (δ) are expressed in parts per million (ppm) and are referenced to $CDCl_3$ (7.26 ppm), DMSO (2.50 ppm), MeCN-$d_3$ (1.94 ppm), or Acetone-$d_6$ (2.05 ppm). Coupling constants are reported as Hertz (Hz). Splitting patterns are indicated as follows: s, singlet; d, doublet; t, triplet; q, quartet; qq, quartet of quartets; dd, doublet of doublet; p, pentet; heptet, heptet; m, multiplet; br, broad singlet. High resolution mass spectra (ESI EI) were measured by the QB3/Chemistry mass spectrometry service at University of California, Berkeley. High performance liquid chromatography (HPLC) and low-resolution ESI Mass Spectrometry were performed on an Agilent Infinity 1200 analytical instrument coupled to an Advion CMS-L ESI mass spectrometer. The column used was Phenomenex Luna 5 µm C18(2) (4.6 mm I.D. × 150 mm) with a flow rate of 1.0 mL/min. The mobile phase was MilliQ-$H_2O$ with 0.05% trifluoroacetic acid (TFA) (eluent A) and HPLC grade MeCN with 0.05% TFA (eluent B). Signals were monitored at 210, 254, 350, 480 and 520 nm over 13 min, with a gradient of 5 to 100% eluent B for 10 min, then held at 100% B for 3 min. Preparative HPLC method was run on an Agilent Technologies 1260 Infinity system, using a Phenomenex Luna 5 µm C18(2) (10 mm I.D. x 150 mm) with a flow rate of 20.0 mL/min. Mobile phase used was MilliQ-$H_2O$ with 0.05% TFA (eluent A) and HPLC grade MeCN with 0.05% TFA (eluent B). Signals were monitored at 254, 625, and 650 nm over 30 min. Solvent was held at 30% elutent B from 0 to 1 min, ran at a linear gradient of 30% eluent B to 100% eluent B from 1 to 28 min, was held at 100% eluent B from 28 to 29 min, and then decreased back to 30% eluent B from 29 to 30 min. Loading solutions were ~10 to 20 mg/mL compound in 30% MeCN in MQ-$H_2O$ with 0.05% TFA.

### Spectroscopic Studies

UV-Vis absorbance and fluorescence spectra were recorded using a 2501 Spectrophotometer (Shimadzu) and a Quantamaster 4 L-format scanning spectrofluorometer (Photon Technologies International). The fluorometer is equipped with an LPS-220B 75-W xenon lamp and power supply, A-1010B lamp housing with integrated igniter, switchable 814 photon-counting/analog photomultiplier detection unit, and MD5020 motor driver. Samples were measured in 1-cm path length quartz cuvettes (Starna Cells). The maximum absorption wavelength ($\lambda_{max}$), maximum emission wavelength ($\lambda_{em}$), extinction coefficients (ε), and quantum yields (Φ) were taken in either 1X dPBS pH 7.2, 1X dPBS with 0.1% Sodium Dodecyl Sulfate (SDS), or trifluoroethanol (TFE) with 0.1% trifluoroacetic acid (TFA).

*HaloTag binding*

These studies were performed in a sub-micro 50 µL quartz fluorometer cell. In a separate glass vial a solution was prepared using PBS buffer and the dye stock in DMSO (maximum 0.5% DMSO concentration in solution). When measuring the background absorbance, nothing else was added to the solution. When measuring the absorbance when bound to HaloTag, we added the purified HaloTag protein (2 equivalents relative to the dye) to the solution and incubated for a minimum of 15 minutes before measuring the absorbance. The final volume of the total solution was always 100 µL, with or without the HaloTag protein. When transferring the solution to the cuvette, we used 70 µL of the solution; less than 65 µL could result in a meniscus in the cuvette that would alter the baseline absorbance. The absorbance values of the bound and unbound dye were recorded in triplicate and averaged in order to record the absorbance and fluorescence turn-on. The quantum yields (Φ) of the HaloTag complexes were measured *via* the same method as the other dyes (Values for absorbance and fluorescence turn-on reported in **Table S1**).

### Cell Culture

*Cell culture and transient transfections (HEK cells, Figure 1)*

Human embryonic kidney 293T (HEK293T) cells were passaged and plated onto 12 mm #1.5 glass coverslips pre-coated with Poly-D-Lysine (PDL; 0.1 mg/ml; in 10 mM $Na_3BO_3$; Sigma-Aldrich) to provide a density of ~44,000 cells/cm$^2$. HEK293T cells were plated and maintained in Dulbecco's modified eagle medium (DMEM) supplemented with 4.5 g/L D-glucose (high glucose), 10% FBS and 1% Glutamax. Unless otherwise stated, all cell culture for imaging experiments were done in high glucose DMEM. For loading cells, dyes were diluted in DMSO to 1000× the final indicated concentration and then diluted 1:1000 in HBSS.

Transfections were done using Lipofectamine 3000 (Invitrogen) 24 h after initial plating. To each well containing a 12 mm coverslip in a 24-well plate, 500 ng DNA/lipofectamine solutions per coverslip were added. The cells were then left untouched for another 24 hours (~48 hours after initial plating) to result in a final confluency of ~175,000 cells/cm$^2$ (~75%) for epifluorescence imaging. Non-transfected cells were plated at the same time as transfected cells, but the addition of DNA/lipofectamine solutions was omitted.

Transfections for electrophysiology experiments began with plating cells at a density of ~61,000 cells/cm$^2$ in a 6-well plate. After 24 hours, cells were transfected with the same lipofectamine reagents at a concentration of 1000 ng DNA/lipofectamine solution per well of the 6-well plate. The cells were then allowed to grow for another 24 hours before being plated onto 25 mm #1.5 glass coverslips precoated with PDL at a density of 26,000 cells/cm$^2$ in DMEM supplemented with 1 g/L D-glucose (low glucose), 10% FBS and 1% Glutamax to achieve single cell confluency. These cells were then allowed to grow for a final 24 hours before imaging and patch clamp electrophysiology experiments.

*Cell culture and transfection (COS-7 cells, Figure 2).*

COS-7 cells were maintained in Dulbecco's Modified Eagle's Medium (DMEM) with 10% fetal bovine serum (FBS) and 1× non-essential amino acids (NEAA). Cells for imaging were seeded onto 18 mm #1.5 coverslips at a density of ~40,000/cm$^2$. 24-48 hours before imaging, cells were transfected with the plasmids described below using Lipofectamine 3000 Transfection Reagent (Thermo Fisher) according to the manufacturer's protocol. HaloTag-Sec61b and ER-HaloTag (HaloTag-ER-3) were described in Wang, *et al.*[1] (Addgene Plasmids #186959 and #186960). ER-Dendra2 (Dendra2-ER-5) was a gift from Michael Davidson (Addgene plasmid #57716). TOM20-mEos4b (mEos4b-TOMM20-N-10) was a gift from Michael Davidson and Loren Looger (Addgene plasmid #57518). Vimentin-HaloTag was constructed through the Gibson assembly (New England BioLabs E2611) of pcDNA3.1(+) backbone, PCR-amplified Halotag from HaloTag-ER-3, and PCR-amplified Vimentin from mCherry-Vimentin-7 (a gift from Michael Davidson, Addgene #55156). Plasmids were amplified in XL1-Blue cells and extracted with the QIAprep Spin Miniprep Kit (QIAGEN). DNA sequences were verified by Sanger sequencing at the UC Berkeley DNA Sequencing Facility.

*Neuron culture*

All animal procedures were approved by the UC Berkeley Animal Care and Use Committees and conformed to the NIH Guide for the Care and Use and Laboratory Animals and the Public Health Policy.

Hippocampi were dissected from embryonic day 19 Sprague Dawley rats (Charles River Laboratory) in cold HBSS (-$Ca^{2+}$, -$Mg^{2+}$, phenol red). Dissected hippocampal tissue was treated with trypsin (2.5%) for 20 min at 37 °C; hippocampi were then triturated with flame-polished Pasteur pipettes in minimum essential media (MEM) supplemented with 5% FBS, 1% GlutaMax, 2% B-27, and 1% 1M dextrose (Fisher Scientific). Dissociated cells were plated onto 25 mm diameter glass coverslips (Fisher Scientific) pre-treated with PDL (as above) at a density of 100,000 cells per coverslip in MEM supplemented media. Neurons were maintained in a humidified incubator at 37 °C with 5% $CO_2$. After 1 day in vitro (DIV), half of MEM supplemented media was removed and replaced with Neurobasal media (NB) supplemented with 2% B-27 and 1% GlutaMax. Evoked and spontaneous imaging was performed on mature neurons 12-13 DIV.

**Fluorescence microscopy**

*Epifluorescence imaging (Figure 1)*

Epifluorescence imaging was performed on an AxioExaminer Z-1 (Zeiss) equipped with a Spectra-X Light engine LED light (Lumencor), controlled with Slidebook (v6, Intelligent Imaging Innovations). Images were acquired with a W-Plan-Apo 63x/1.0 water objective (63x; Zeiss). Images were focused onto an OrcaFlash4.0 sCMOS camera (sCMOS; Hamamatsu). More detailed imaging information for each experimental application is expanded below.

Transfected and non-transfected HEK293T cells were co-stained with 1 µg/mL (1.7 µM) Hoechst 33342 and 500 nM HaloTag ligand (**BF$_{646\text{-}xHTL}$**) in HBSS for 20 min at 37°C, 5% $CO_2$. Coverslips were then washed with 1 volume of HBSS before being transferred to a 3 cm imaging dish containing 3 mL HBSS ("wash conditions", Figure S1). For "no wash" conditions, imaging was performed in the staining solution (Figure S1). Images were acquired with a W-Plan-Apo 63x/1.0 water objective (Zeiss) and OrcaFlash4.0 sCMOS camera (Hamamatsu). For HaloTag ligand imaging, the excitation light was delivered from an LED with a power of 0.094 W/cm$^2$ (ND: 75) for 2 ms exposures at 631/28 nm (bandpass) and emission was collected through a quadruple emission filter (430/32, 508/14, 586/30, 708/98 nm) after passing through a quadruple dichroic mirror (432/38, 509/22, 586/40, 654 nm LP). For Hoechst 33342 imaging, the excitation light was delivered from an LED with a power of 0.018 W/cm$^2$ (ND: 75) for 100 ms exposures at 390/22 nm (bandpass) and emission was collected with the same quadruple emission filter. Finally, a DIC image was also taken for each field of view (FOV). Images were then exported as tiff files to be analyzed in ImageJ.

*Imaging sample preparation (Figure 2)*

Live-cell imaging was performed in Leibovitz's L-15 medium (ThermoFisher, 21083027) supplemented with 20 mM HEPES (pH 7.5) and 2.5-nM BF-646-xHTL(T5) or SiR-xHTL(T5). Cells were incubated with the above dye-containing medium for 20–30 min and then imaged directly without washing. For fixed-cell imaging, cells were fixed with 3% paraformaldehyde (Electron Microscopy Sciences, 15714) and 0.1% glutaraldehyde (Electron Microscopy Sciences, 16365) in DPBS at room temperature for 20 min, and then washed twice with a freshly prepared 0.1% (wt/wt) $NaBH_4$ solution followed by three additional washes with DPBS. The fixed cells were subsequently incubated with DPBS containing 2.5-nM BF-646-xHTL(T5) and imaged in this medium.

*PAINT, SMdM, and PALM super-resolution microscopy (Figure 2)*

Single-molecule imaging was performed on a homebuilt inverted microscope[2] using a Nikon CFI Plan Apo λ 100x oil-immersion objective (NA = 1.45). A 647-nm laser excited the sample at ~1.5 kW/cm$^2$, which entered the sample slightly below the critical angle of total internal reflection to illuminate a few micrometers into the sample. For PAINT[3], the reversible binding of BF-646-xHTL(T5) or SiR-xHTL(T5) to the expressed HaloTag in the cell yielded on-off blinking of single-molecule fluorescence, which was recorded continuously using an Andor iXon Ultra 897 EM-CCD camera at 56 or 109 fps. ~60,000 frames were typically recorded per image. For 3D localization, a cylindrical lens was inserted to introduce astigmatism and encode single-molecule depth (Z) information.[5] Single-molecule images in all frames were localized and drift-corrected as described previously.[4-6] PAINT images were rendered by temporally segmenting the resulting single-molecule localizations by frame, with ~5,000 frames typically providing good coverage of cellular structures, yielding ~1-min temporal resolution. SM*d*M was performed similarly as above, but with the excitation modulated as tandem 1-ms duration pulses across paired frames[7-8] at a center-to-center separation of $\Delta t$ = 2 ms. Vectoral displacements were extracted between the paired frames and spatially binned with a 100-nm grid, and the displacements in each spatial bin were analyzed to determine the local principal direction of diffusion and diffusion coefficient.[9] Concurrent two-color live-cell PALM-PAINT imaging was performed by alternating the excitation laser between 561 nm (for mEos4b) and 647 nm (for BF-646-xHTL(T5)) between frames. Weak (~0.01 W/cm$^2$) 405 nm illumination was further applied to photoconvert a small fraction of mEos4b to the 561 nm-excitable state and maintain the count of single molecules in the 561 nm channel. Concurrent two-color SM*d*M was performed by alternating between tandem 561-nm (for Dendra2) and 647 nm (for BF-646-xHTL(T5)) excitation pairs.

**DNA Constructs / Molecular Biology**

**Table S1.** Plasmids

| Target | Plasmid | Reference / Sequence |
|---|---|---|
| Nucleus | pTG735-3xFlag-Halo-3xNLS | https://benchling.com/s/seq-dfPcBKdXECGwrhPqD6P9?m=slm-3UY7EYrK9TMwpKXFWjuM |
| ER membrane | HaloTag-sec61b | Addgene #186959 |
| ER lumen | ER-HaloTag (HaloTag-ER-3) | Addgene #186960 |
| ER lumen | ER-Dendra2 (Dendra2-ER-5) | Addgene #57716 |
| Mitochondria outer membrane | mEos4b-TOMM20-N-10 | Addgene #57518 |
| Vimentin | pcDNA3.1 (+) vimentin-HaloTag | https://benchling.com/s/seq-g1QZQBTyAn1JiZO6zXqE?m=slm-1ScNFtkCLxOcm56dOfY0 |

Dendra2-ER-5 was a gift from Michael Davidson (Addgene plasmid # 57716 ; http://n2t.net/addgene:57716 ; RRID:Addgene_57716)

mEos4b-TOMM20-N-10 was a gift from Michael Davidson & Loren Looger (Addgene plasmid # 57518 ; http://n2t.net/addgene:57518 ; RRID:Addgene_57518)

**Supporting Schemes**

**Scheme S1.** Synthesis of fluorescent xHTL ligands

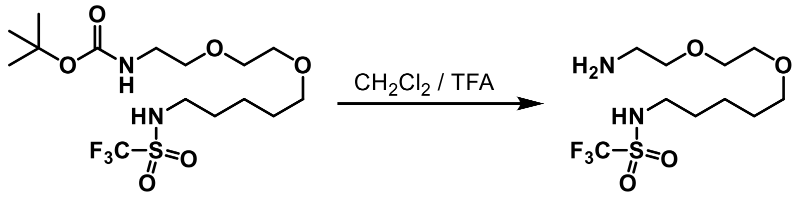

SI-1 SI-2

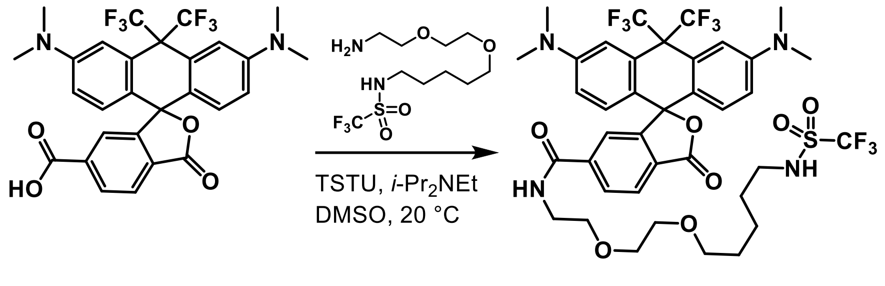

BF-646 BF-646-xHTL(T5)

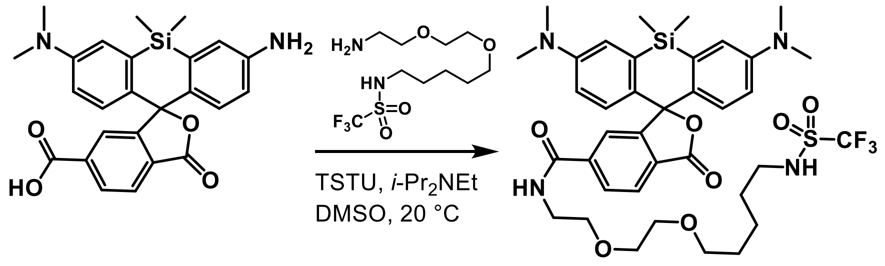

SiR SiR-xHTL(T5)

### Detailed Synthetic Procedures

*Amine SI-2*

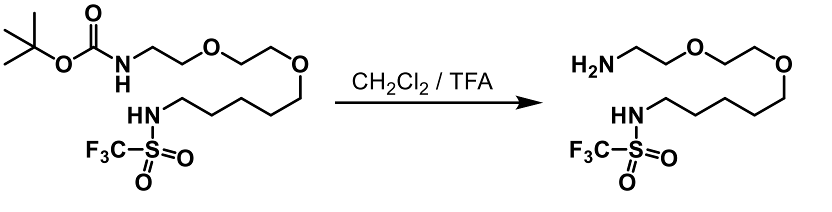

A vial was charged with the Boc-protected amine **SI-1** (synthesized according to Kompa, *et al.*)[10] (34.0 mg, 0.0805 mmol, 1.0 equiv.) before adding $CH_2Cl_2$ (0.75 mL, 0.11 M of **SI-1**) and trifluoroacetic acid (0.75 mL, 0.11 M of **SI-1**). After 30 minutes, toluene (1.0 mL) was added to the reaction, and the solvent was removed *in vacuo*. After fully removing the solvent, a second addition of toluene (1.0 mL) was made. The vial was sonicated before again removing the solvent *in vacuo*. The resulting crude material was placed under vacuum for 12 hours before dissolving in DMSO (1.68 mL) to create a 0.048 M stock solution of amine **SI-2**.

*BF646-xHTL(T5) (**1**)*

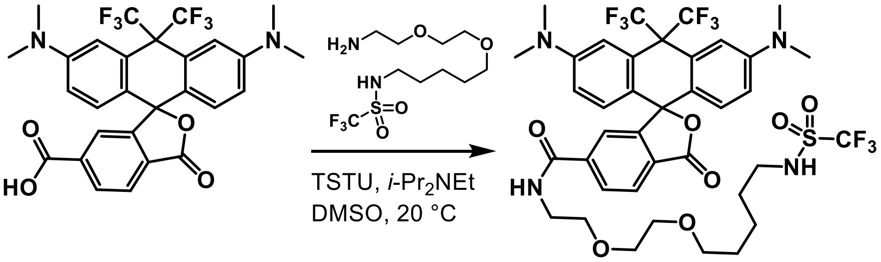

A vial with **BF646** (9.6 mg, 0.017 mmol, 1.0 equiv.) and a stir bar was charged with DMSO (0.49 mL, 0.016 M of **BF646**). Diisopropylethylamine (0.03 mL, 0.17 mmol, 10 equiv.) was added, creating a transparent solution. To this transparent solution was added TSTU (6.5 mg, 0.022 mmol, 1.3 equiv.). After 20 minutes, a solution of **SI-2** in DMSO (0.57 mL, 0.0272 mmol, 1.6 equiv.) was added. Monitoring the reaction by analytical LCMS indicated that the reaction was complete within 50 minutes. The reaction was quenched by diluting with a $H_2O$/MeCN mix and directly purifying this solution on the preparative HPLC (10 to 100% MeCN/$H_2O$ with 0.05% trifluoroacetic acid). The solvent was removed *in vacuo* from the fractions containing **BF646-xHTL(T5)** before dissolving the material in MeOH and filtering through a C-18 silica pipette column. The solvent was removed *in vacuo* before dissolving in MeOH and filtering through a second C-18 silica pipette column. Removal of the solvent yielded **BF646-xHTL(T5)** (11.5 mg, 0.013 mmol, 78% yield) as a transparent film.

Note: As we previously reported, the bis-(trifluoromethyl)methylene group requires special consideration when performing NMR experiments. The advantage of this functional group is the ability to quickly verify the purity of your compound when accessing the presence of additional biproducts containing the bis-(trifluoromethyl)methylene group when examining the NMR spectra. In **BF646-xHTL(T5)** (**1**) there exists two signals in the $^{19}F$ NMR due to the diastereotopic trifluoromethyl carbons. There may also be a third signal due to a trifluoroacetate counter anion, depending on what conditions were used to isolate the dye. Since the $^{19}F$ NMR is extremely quick to acquire and very sensitive, it is very useful for assessing sample purity. The downsides of this functional group are that the coupling of $^{19}F$ and $^{13}C$ nuclei results in carbon signals containing multiplicity, significantly decreasing their intensity in the NMR spectra. Even with a highly sensitive NMR instrument containing a cryoprobe, we often found it challenging to identify the peaks corresponding to the trifluoromethyl carbons and the methylene carbon bearing the trifluoromethyl groups. Even with these advantages, we often found it necessary to perform $^{19}F$ decoupled $^{13}C$ NMR experiments in order to identify these signals.

$^1H$ NMR (600 MHz, $CDCl_3$) δ 8.04 (dd, $J$ = 8.0, 0.8 Hz, 1H), 7.92 (dd, $J$ = 8.0, 1.5 Hz, 1H), 7.35 (s, 2H), 7.21 – 7.19 (m, 1H), 6.75 – 6.71 (m, 4H), 6.62 – 6.59 (m, 1H), 3.63 – 3.60 (m, 2H), 3.59 – 3.57 (m, 2H), 3.55 (q, $J$ = 5.3 Hz, 2H), 3.49 –

3.46 (m, 2H), 3.32 (t, *J* = 5.9 Hz, 2H), 3.21 (t, *J* = 6.6 Hz, 2H), 3.00 (s, 12H), 1.52 (p, *J* = 6.9 Hz, 2H), 1.47 (p, *J* = 6.4 Hz, 2H), 1.33 (p, *J* = 7.2 Hz, 2H).

$^{13}$C NMR (151 MHz, $CDCl_3$) δ 170.1, 166.5, 156.7, 150.3, 141.2, 129.7, 128.3, 128.2, 126.4, 125.5, 121.8, 121.0, 115.0, 113.2, 86.0, 70.9, 70.4, 70.2, 69.6, 44.1, 40.3, 40.1, 28.6, 23.0.

$^{13}$C{$^{19}$F} NMR (151 MHz, $CDCl_3$) δ 124.3, 124.2, 119.9, 55.6.

$^{19}$F NMR (565 MHz, $CDCl_3$) δ -63.5 (q, *J* = 7.2 Hz), -63.8 (q, *J* = 7.0 Hz), -77.4 (s).

HR-ESI-MS m/z for $C_{40}H_{42}O_7N_4F_9S_1^+$ $[M+H]^+$: 869.2630. Calculated: 869.2625

*SiR-xHTL(T5)*

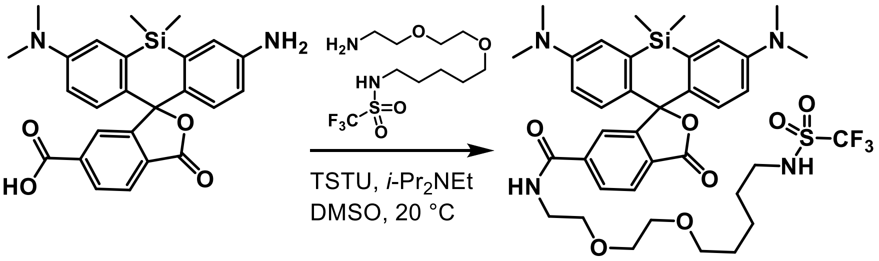

**SiR** **SiR-xHTL(T5)**

A vial with stir bar and **SiR** (7.0 mg, 0.015 mmol, 1.0 equiv.) was charged with DMSO (0.44 mL, 0.016 M of **SiR**) and *i*-$Pr_2Net$ (0.03 mL, 0.17 mmol, 11 equiv.). To this faint green solution was added TSTU (6.0 mg, 0.020 mmol, 1.3 equiv.). After 20 minutes, a solution of amine **SI-2** (0.49 mL, 0.048 M in DMSO, 0.024 mmol, 1.6 equiv.) was added. The reaction was complete by HPLC-MS within an hour. The reaction was quenched with $H_2O$/MeCN and purified by preparative HPLC (10 to 100% MeCN/$H_2O$, 0.05% TFA). The solvent was removed on the fractions containing the product. The residue was filtered twice through C-18 silica pipette columns using MeOH as a solvent. The product **SiR-xHTL(T5)** (5.3 mg, 0.0068 mmol, 46% yield) was isolated as a transparent film.

$^1$H NMR (600 MHz, $CDCl_3$) δ 8.01 (d, *J* = 8.0 Hz, 1H), 7.95 (dd, *J* = 8.0, 1.3 Hz, 1H), 7.68 (s, 1H), 7.00 (d, *J* = 2.9 Hz, 2H), 6.89 (t, *J* = 5.1 Hz, 1H), 6.78 (d, *J* = 8.9 Hz, 2H), 6.58 (dd, *J* = 8.9, 2.9 Hz, 2H), 6.40 (s, 1H), 3.66 (t, *J* = 4.8 Hz, 2H), 3.64 – 3.60 (m, 4H), 3.54 – 3.49 (m, 2H), 3.33 (t, *J* = 6.1 Hz, 2H), 3.21 – 3.14 (m, 2H), 2.97 (s, 12H), 1.49 – 1.38 (m, 4H), 1.32 – 1.27 (m, 2H), 0.66 (s, 3H), 0.60 (s, 3H).

$^{13}$C NMR (151 MHz, $CDCl_3$) δ 170.0, 166.6, 149.5, 139.8, 137.3, 131.8, 129.4, 128.5, 127.9, 126.2, 123.5, 120.0 (q, *J* = 321.1 Hz), 117.3, 113.9, 71.0, 70.3, 70.2, 69.6, 44.1, 40.6, 40.2, 28.6, 22.9, 0.5, -1.2.

$^{19}$F NMR (565 MHz, $CDCl_3$) δ -77.4 (s).

HR-ESI-MS m/z for $C_{27}H_{48}O_7N_4F_3S_1Si_1^+$ $[M+H]^+$: 777.2960. Calculated: 777.2956.

**Supporting Figures**

**Figure S1.** Wash out of fluorescent xHTL dyes in HEK293T cells

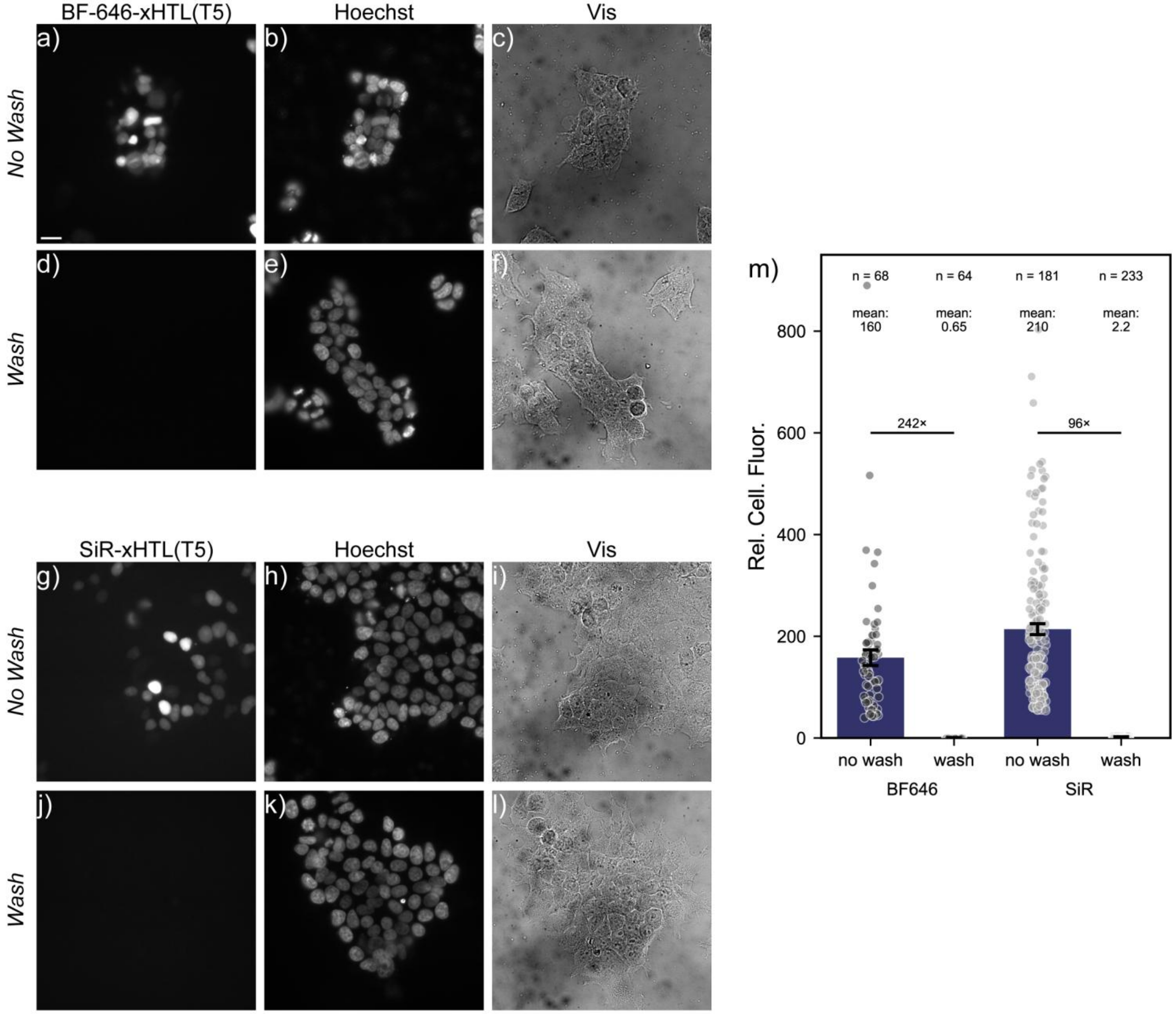

xHTL dyes in wash vs no wash conditions. HEK293T cells expressing nuclear HaloTag (NLS×3-HaloTag) co-stained with **a**) BF-646-xHTL(T5) (200 nM) and **b**) Hoechst imaged directly in staining solution. **c**) Transmitted light image (vis) shown for reference. Cells co-stained with **d**) BF646-xHTL-(T5) (200 nM) and with **e**) Hoechst after wash protocol (removal of the staining solution, replacement with 1× volume of fresh buffer, and transfer to 3 mL of HBSS for imaging). **f**) Transmitted light image (vis) shown for reference. Scale bar 20 µm.

HEK293T cells expressing nuclear HaloTag (NLS×3-HaloTag) co-stained with **g**) SiR-xHTL (200 nM) and **h**) Hoechst imaged directly in staining solution. **i**) Transmitted light image (vis) shown for reference. Cells co-stained with **j**) SiR-xHTL (200 nM) and with **k**) Hoechst after wash protocol. **l**) Transmitted light image (vis) shown for reference. **m**) Plot of nuclear ROIs mean fluorescence intensity of BF-646-xHTL vs SiR-xHTL in *no wash* vs *wash* conditions. Data are mean ± SEM from the indicated number of nuclear regions of interest (ROI). The mean fluorescence intensity is reported for each condition, and the ratio of fluorescence intensity between no wash and wash conditions for each dye is listed (242× for BF-646-xHTL; 96× for SiR-xHTL).

**Figure S2.** Live-cell PAINT with BF-646-xHTL(T5) in ER (membrane) of COS-7 cells

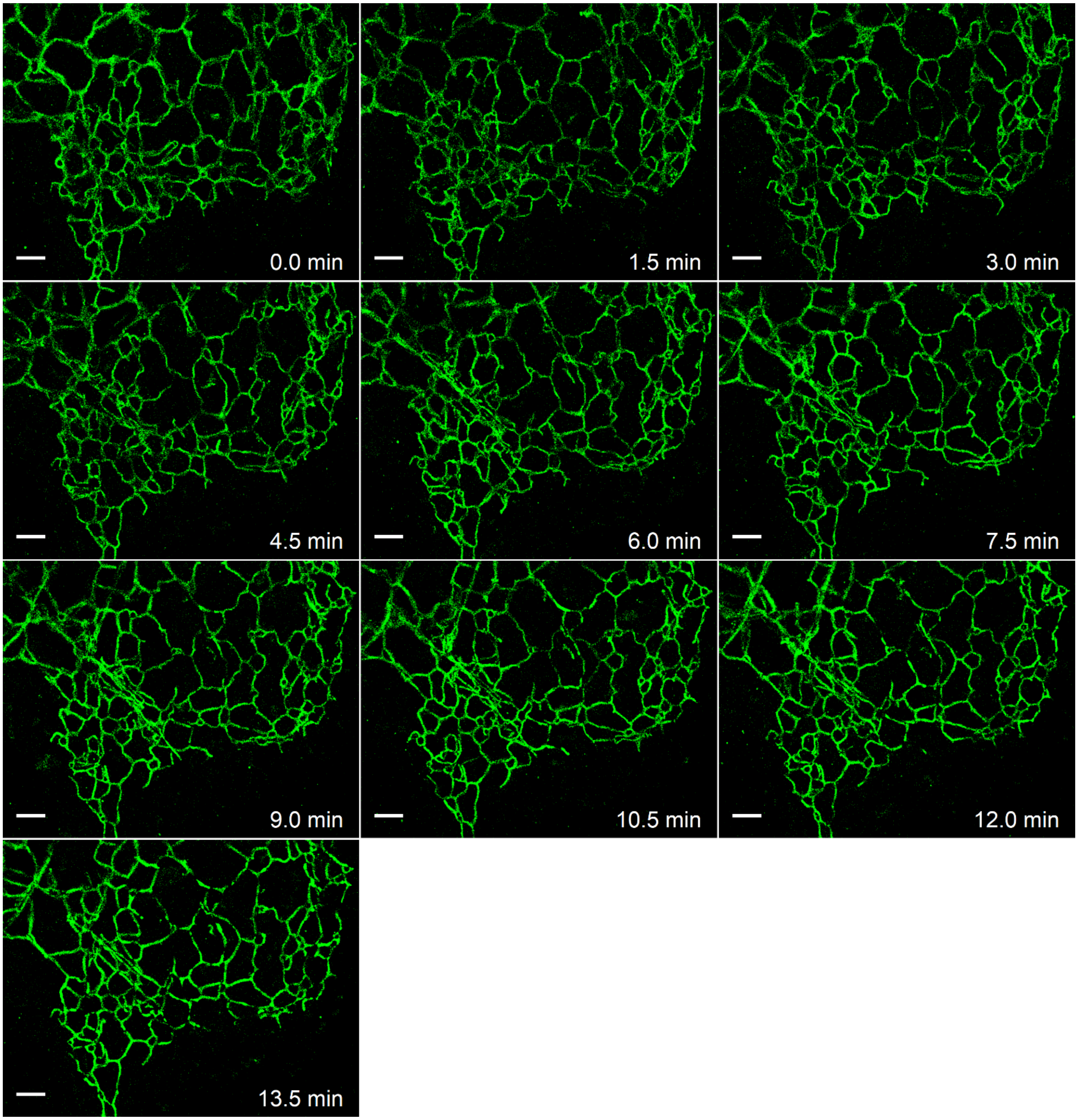

Time series of live-cell PAINT images of BF-646-xHTL(T5)-labeled HaloTag-Sec61β targeting the ER membrane in a living COS-7 cell, constructed with a 1.5-min temporal resolution (5,000 frames at 56 fps). Scale bar: 2 μm.

**Figure S3.** Live-cell PAINT with BF-646-xHTL(T5) in ER (lumen) of COS-7 cells

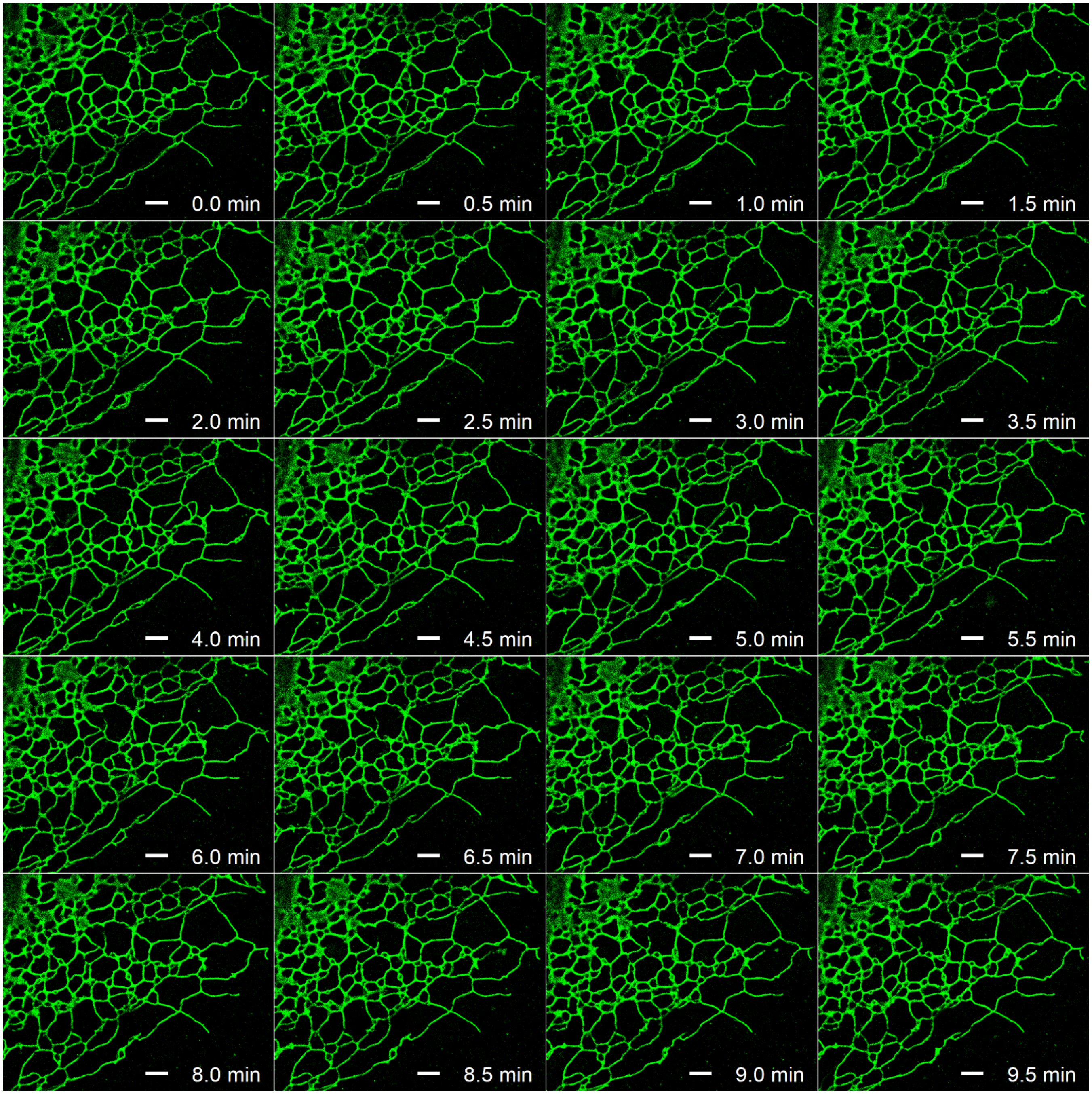

Time series of live-cell PAINT image with BF-646-xHTL(T5) labeling ER-HaloTag diffusing in the ER lumen, constructed with a temporal resolution of 30 s (3,270 frames at 109 fps). Scale bar: 2 µm.

**Figure S4.** Live-cell PAINT image of BF-646-xHTL(T5)-labeled vimentin-HaloTag in a COS-7 cell

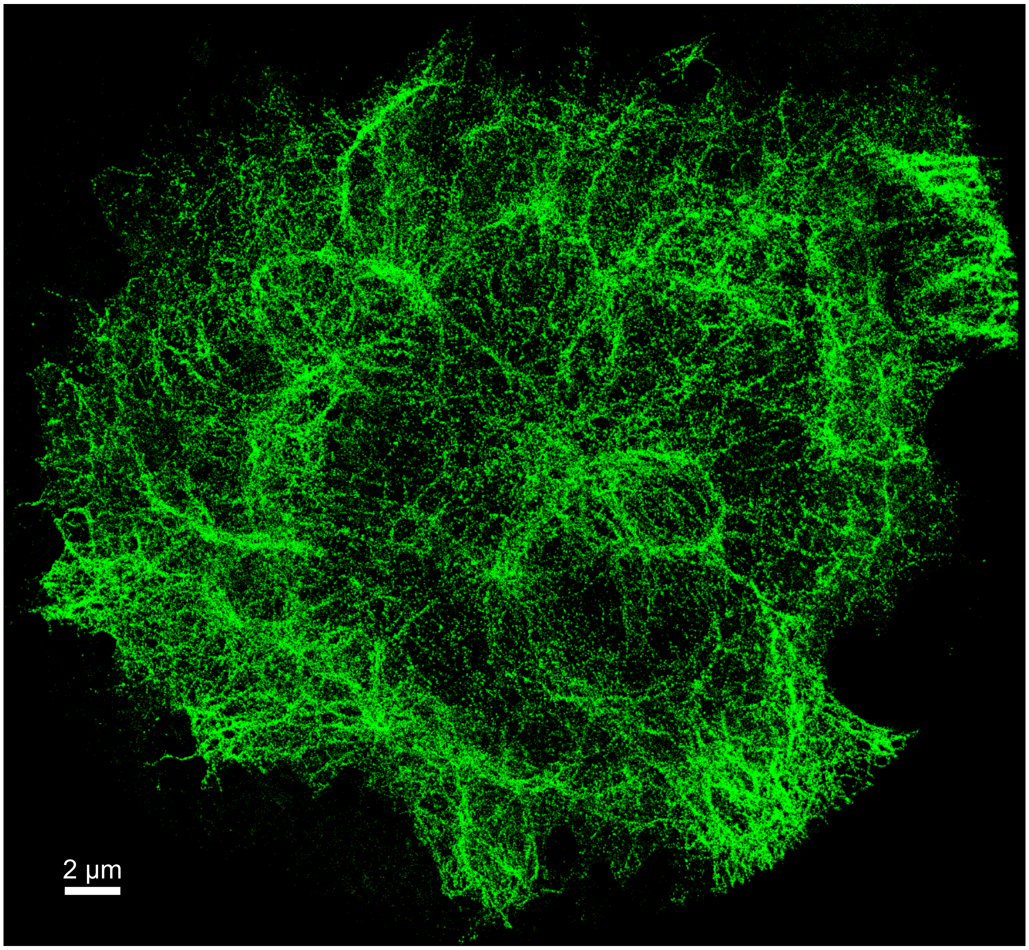

Live-cell PAINT image of BF-646-xHTL(T5)-labeled vimentin-HaloTag in a COS-7 cell.

**Figure S5.** Live-cell PAINT with BF-646-xHTL(T5) in ER (membrane) of neurons

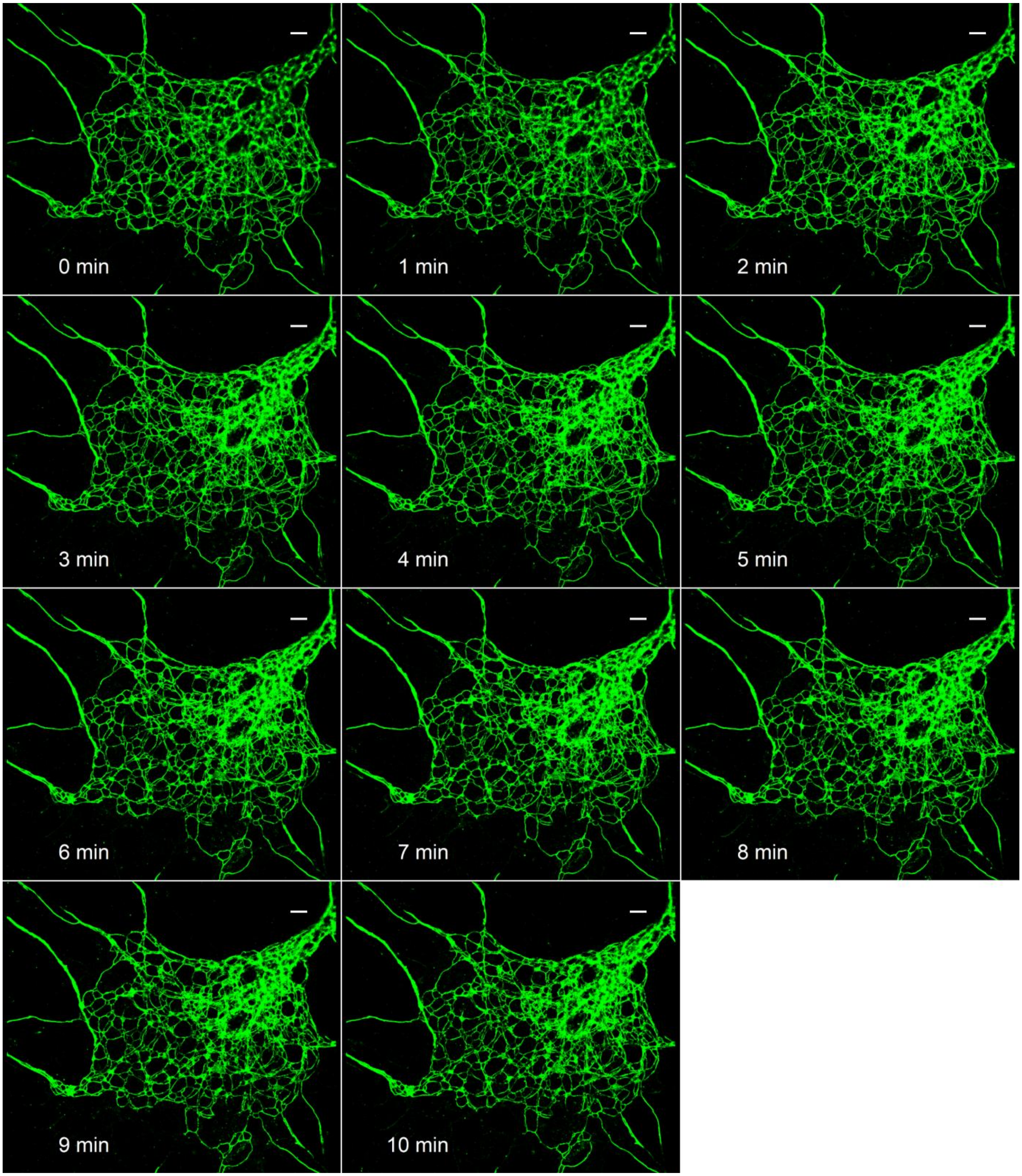

Time series of live-cell PAINT images of BF-646-xHTL(T5)-labeled HaloTag-Sec61β targeting the ER membrane in a cultured primary hippocampal neuron, constructed at a temporal resolution of 1 min (6,504 frames at 108 fps). Scale bar: 2 μm.

**Figure S6.** Time series of two-color live-cell PALM-PAINT

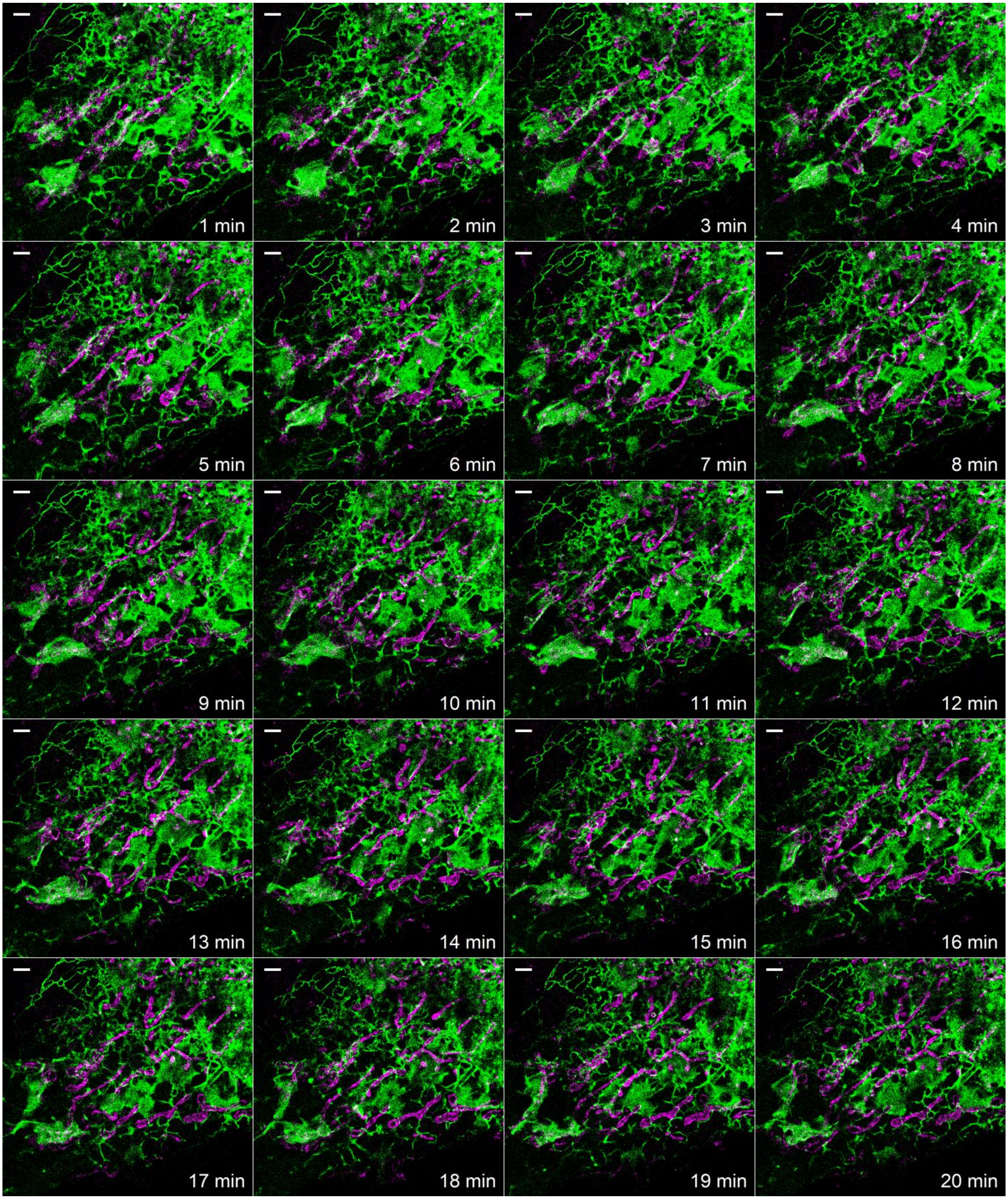

Time series of two-color live-cell PALM-PAINT of TOM20-mEos4b (magenta) and BF-646-xHTL(T5)-labeled HaloTag-Sec61β (green) in a COS-7 cell, constructed at a temporal resolution of 1 min. Scale bar: 2 μm.

**Figure S7.** Time series of SM*d*M super-resolution maps of ER-HaloTag (lumen) labels with BF-646-xHTL(T5)

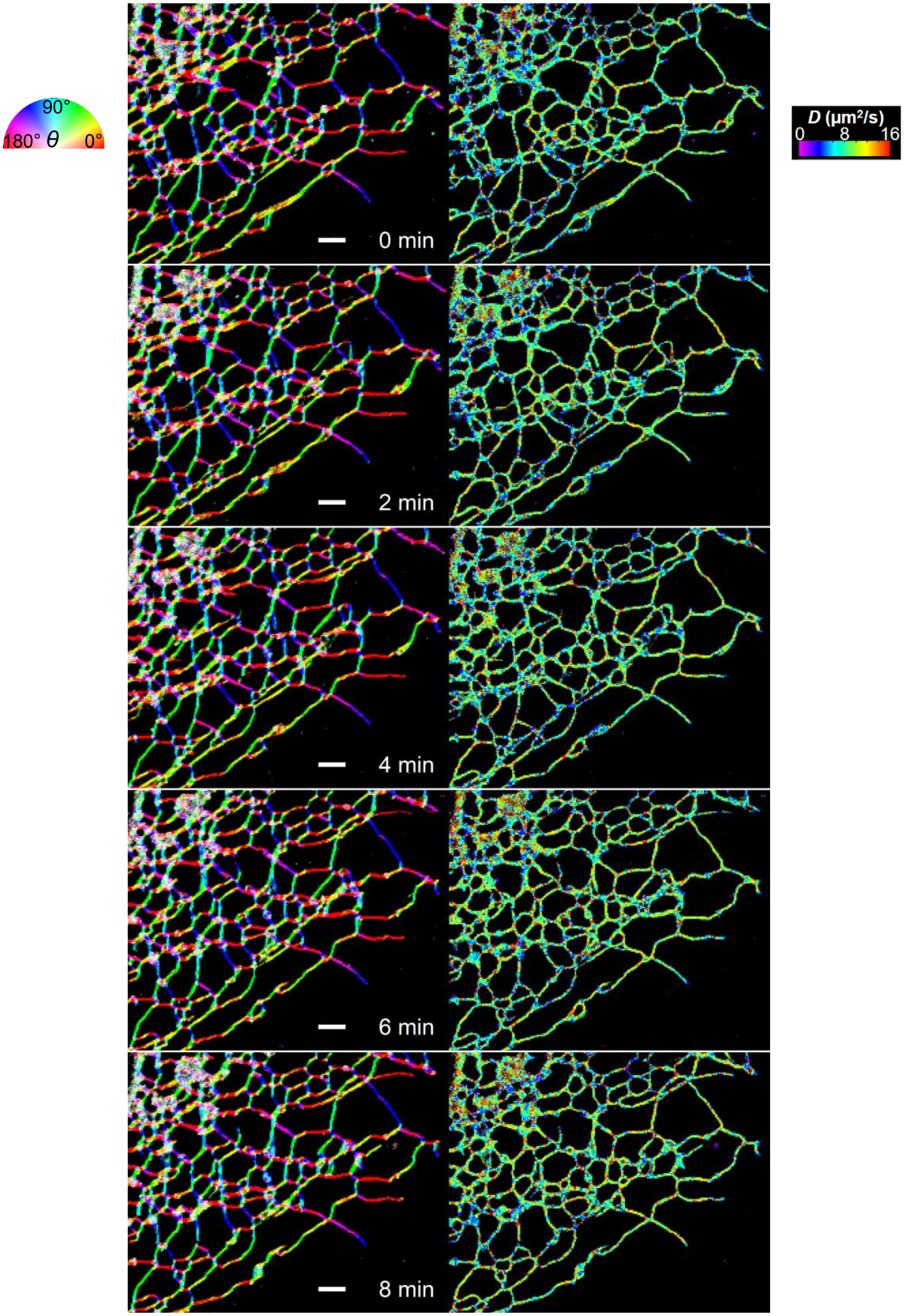

Time series of SM*d*M super-resolution maps of the local principal direction of diffusion (**Left**) and diffusion coefficient (**Right**) for BF-646-xHTL(T5)-labeled ER-HaloTag in a COS-7 cell, constructed at a temporal resolution of 2 min. Scale bar: 2 µm.

### Supplementary Movie Captions

#### Movie S1.

Raw single-molecule data over 50,000 frames for COS-7 cells expressing HaloTag-Sec61β stained with BF-646-xHTL(T5) versus SiR-xHTL(T5), imaged and presented under identical conditions. Single-molecule images were recorded at 56 fps and were shown every 100 frames.

#### Movie S2.

Consecutive time series of live-cell PAINT for a COS-7 cell expressing HaloTag-Sec61β stained with BF-646-xHTL(T5). Scale bar: 2 μm.

#### Movie S3.

Consecutive time series of live-cell PAINT for a COS-7 cell expressing ER-HaloTag stained with BF-646-xHTL(T5). Scale bar: 2 μm.

#### Movie S4.

Consecutive time series of live-cell PAINT for a rat primary hippocampal neuron expressing HaloTag-Sec61β stained with BF-646-xHTL(T5). Scale bar: 2 μm.

#### Movie S5.

Consecutive time series of two-color live-cell PAINT for a COS-7 cell co-expressing HaloTag-Sec61β (green) and mEos4b-TOM20 (magenta). Scale bar: 2 μm.

#### Movie S6.

Consecutive time series of SM*d*M direction map and *D* map for a COS-7 cell expressing ER-HaloTag stained with BF-646-xHTL(T5). Scale bar: 2 μm.

**Supporting Spectra**

**Spectrum S1.** $^{1}$H NMR of BF-646-xHTL(T5)

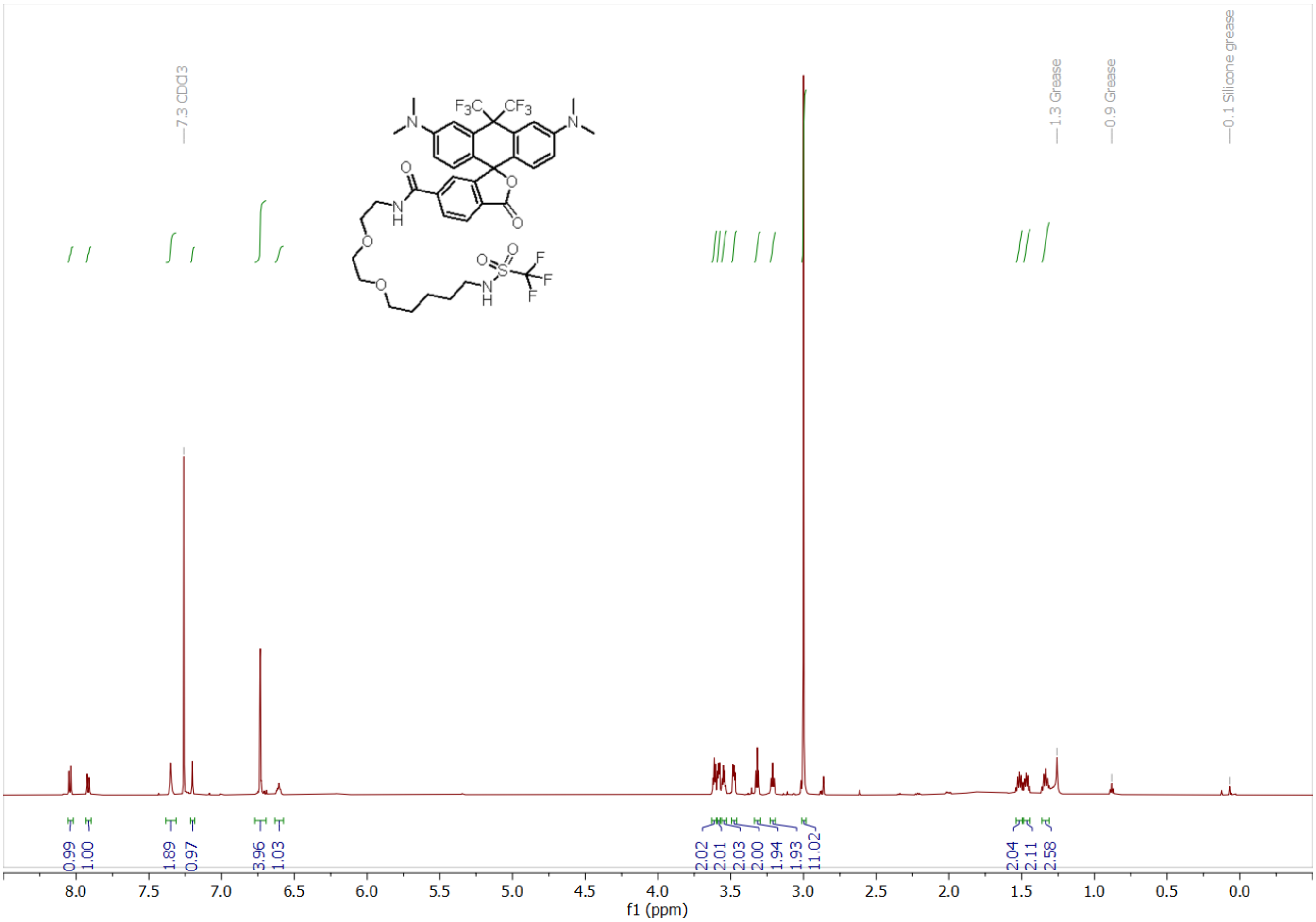

**Spectrum S2.** $^{13}$C NMR of BF-646-xHTL(T5)

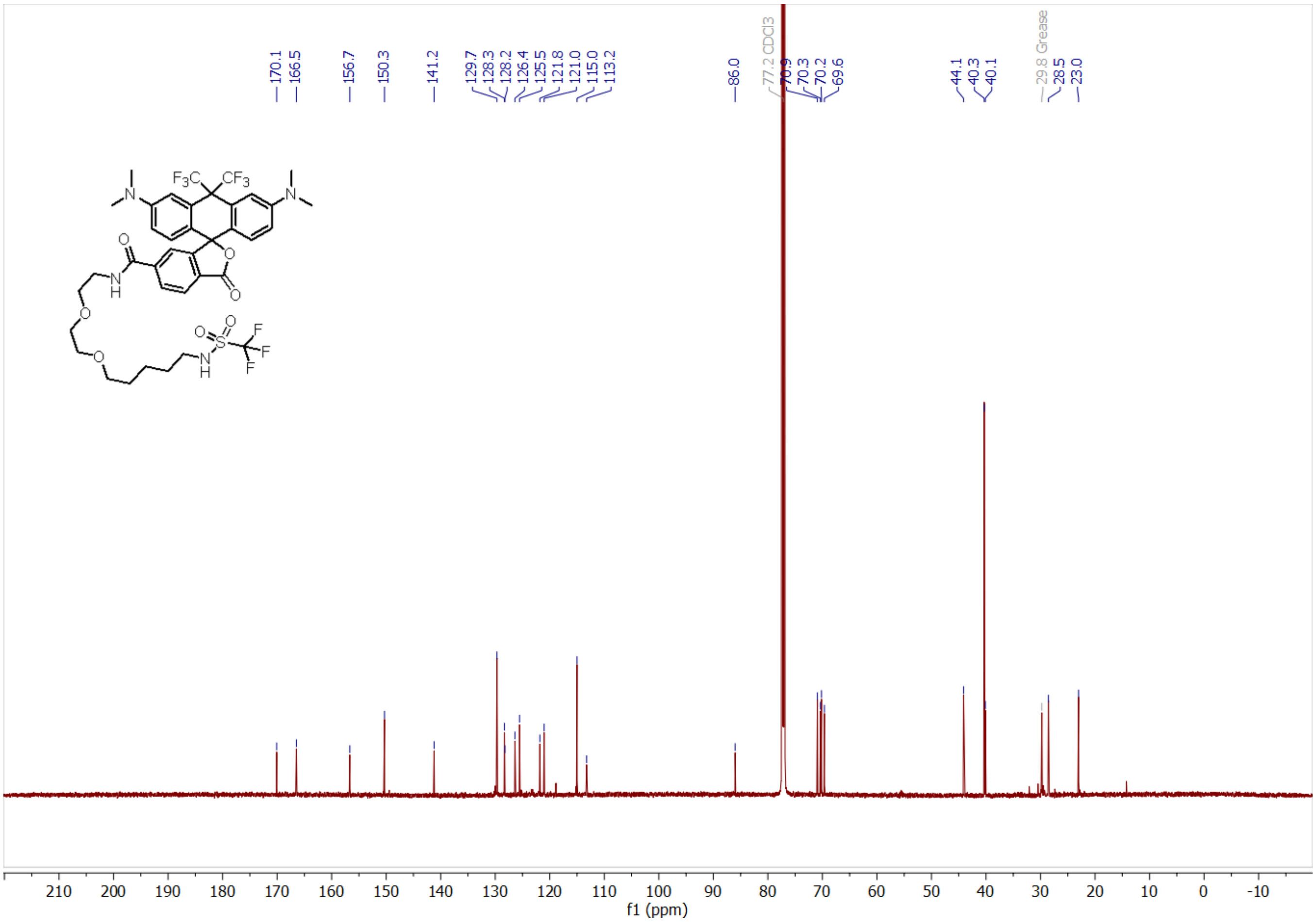

**Spectrum S3.** $^{13}C\{^{19}F\}$ NMR of BF-646-xHTL(T5)

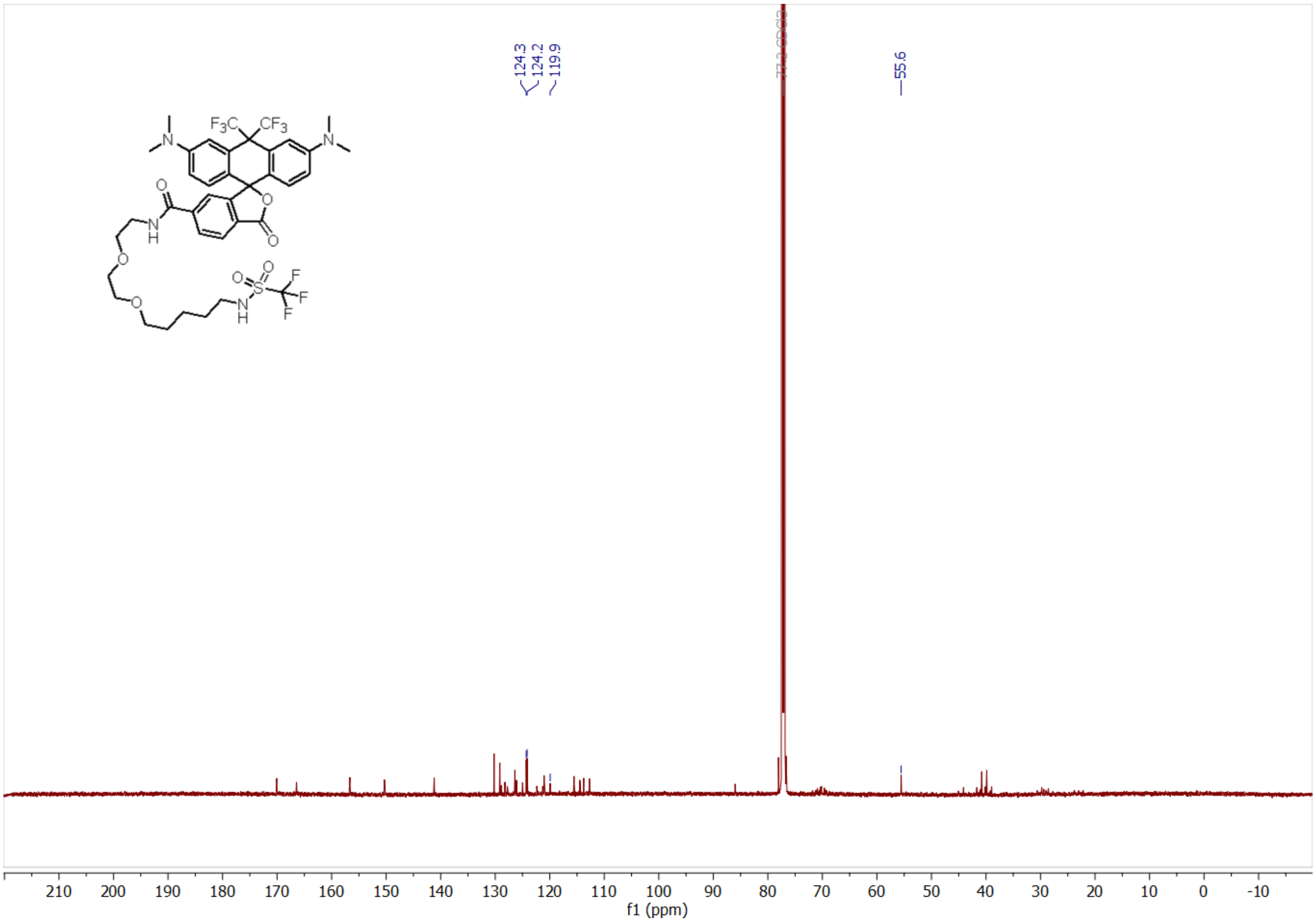

**Spectrum S4.** $^{19}$F NMR of BF-646-xHTL(T5)

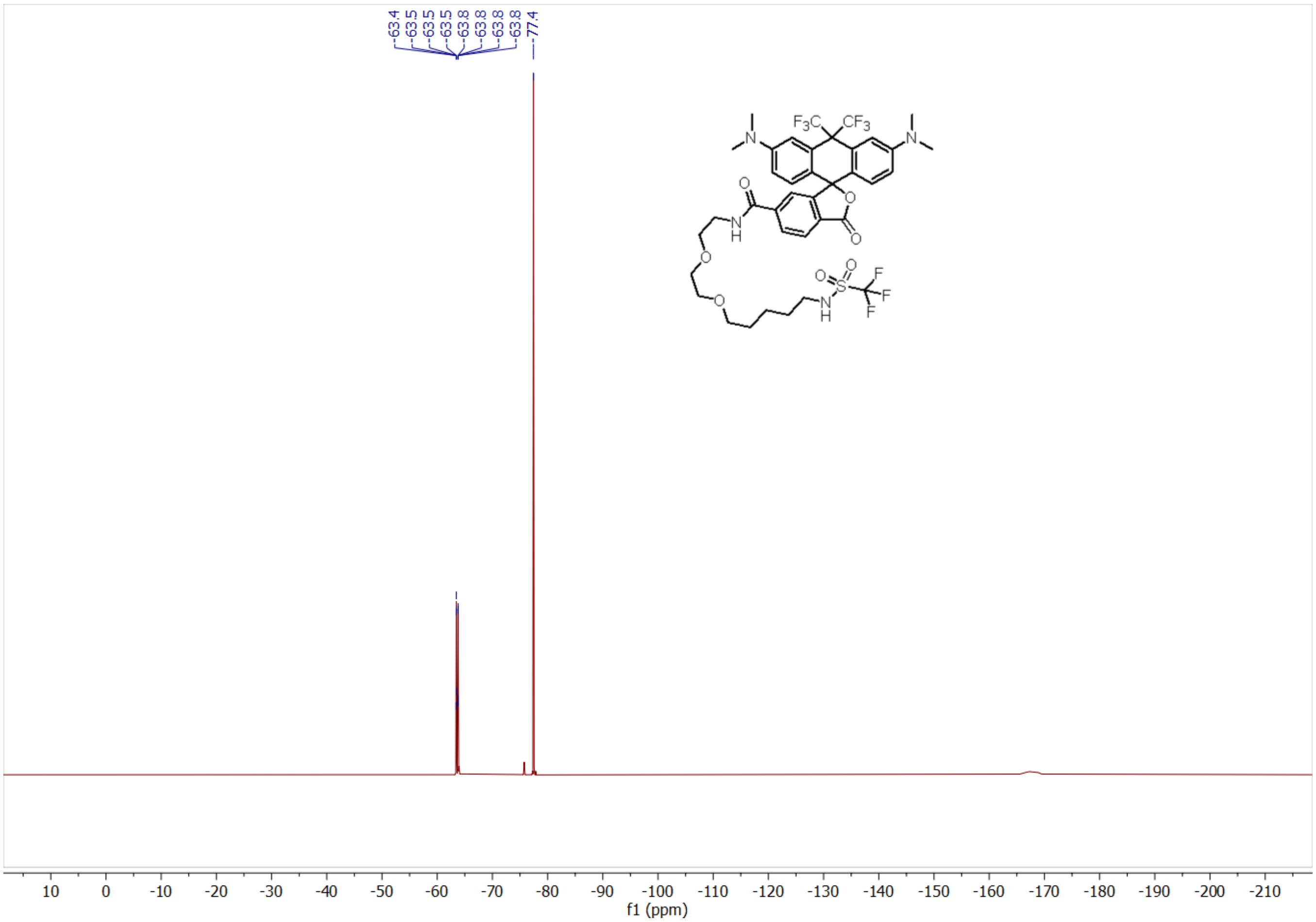

**Spectrum S5.** $^1$H NMR of SiR-xHTL(T5)

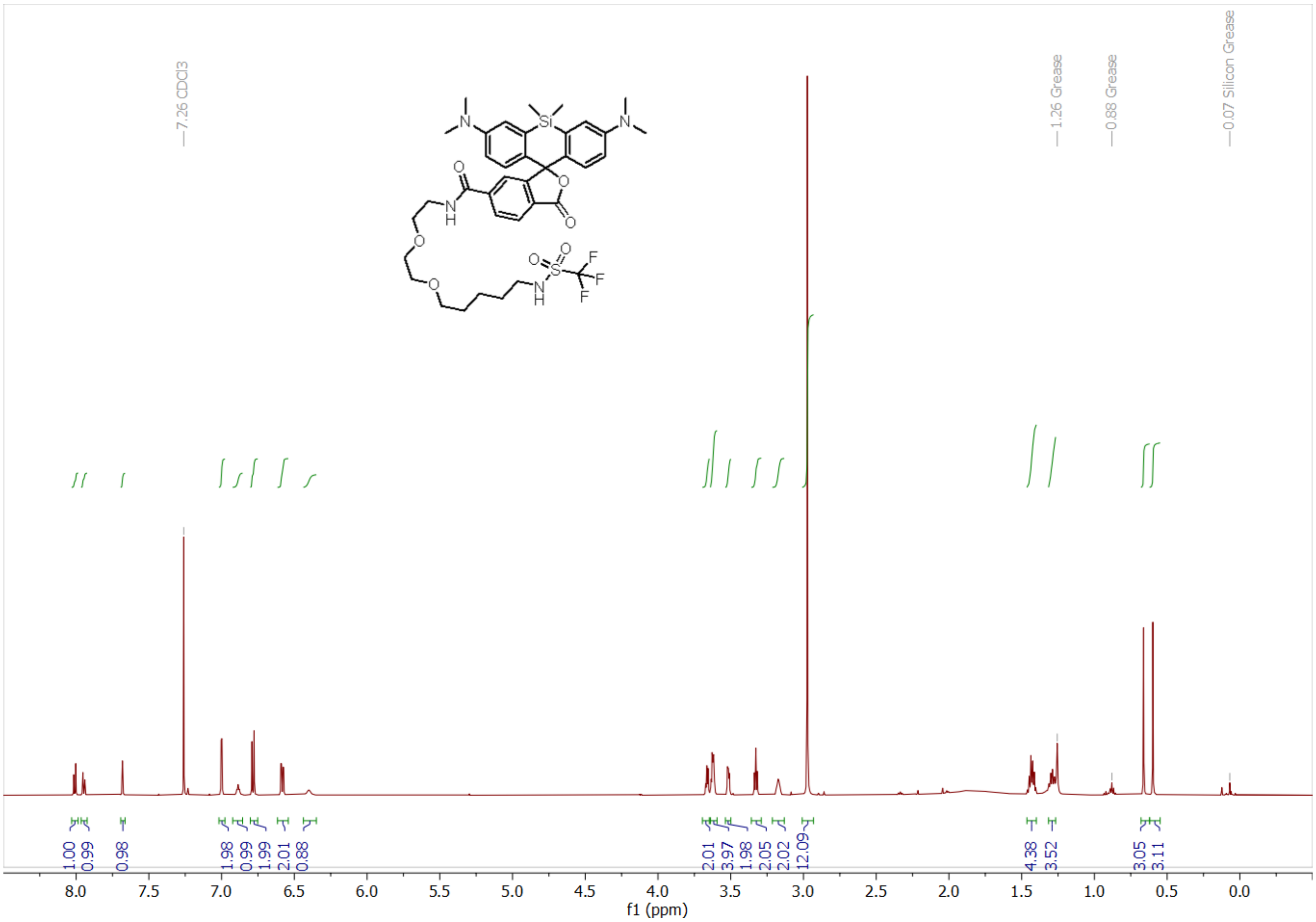

**Spectrum S6.** $^{13}C$ NMR of SiR-xHTL(T5)

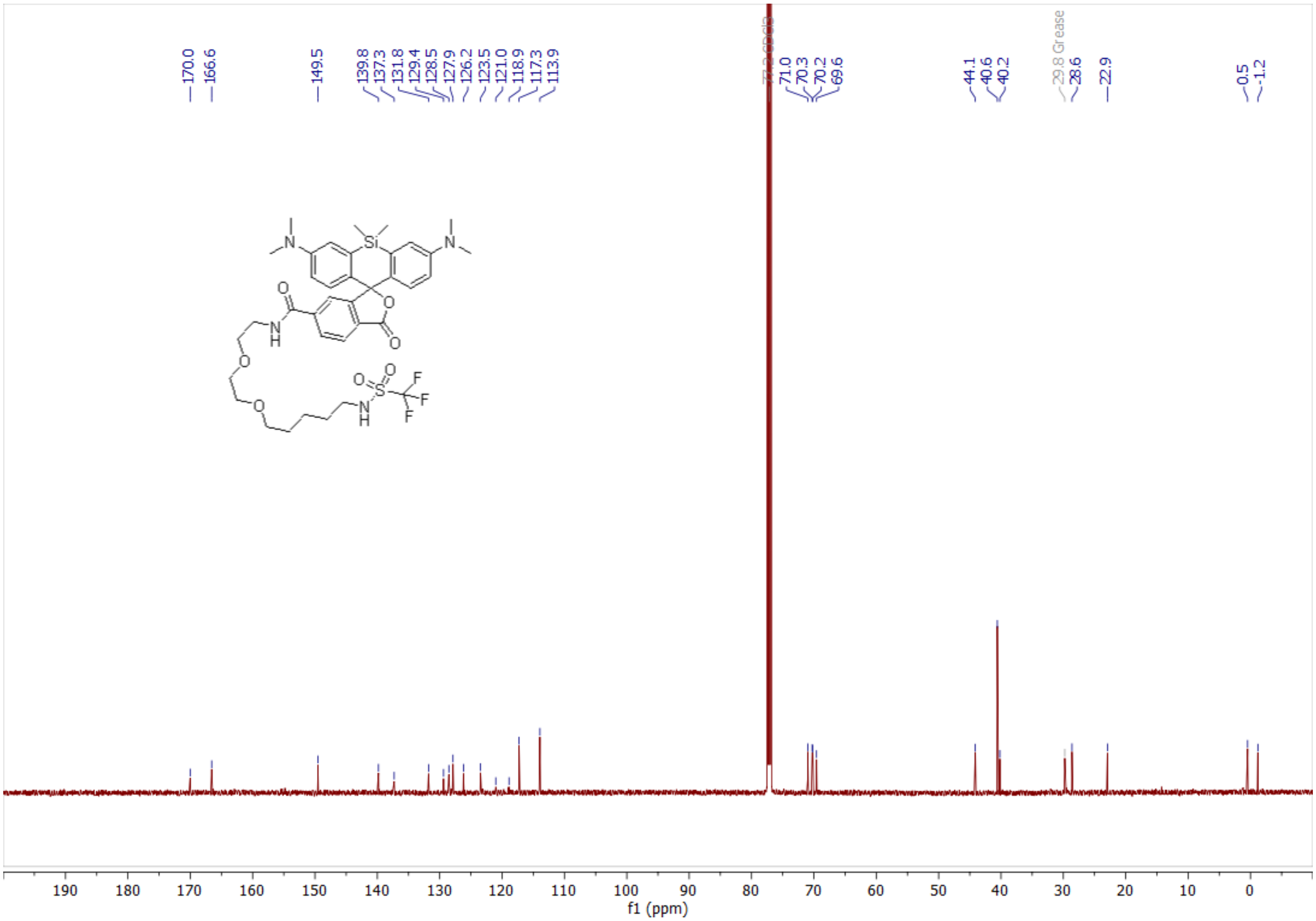

**Spectrum S7.** $^{19}$F NMR of SiR-xHTL(T5)

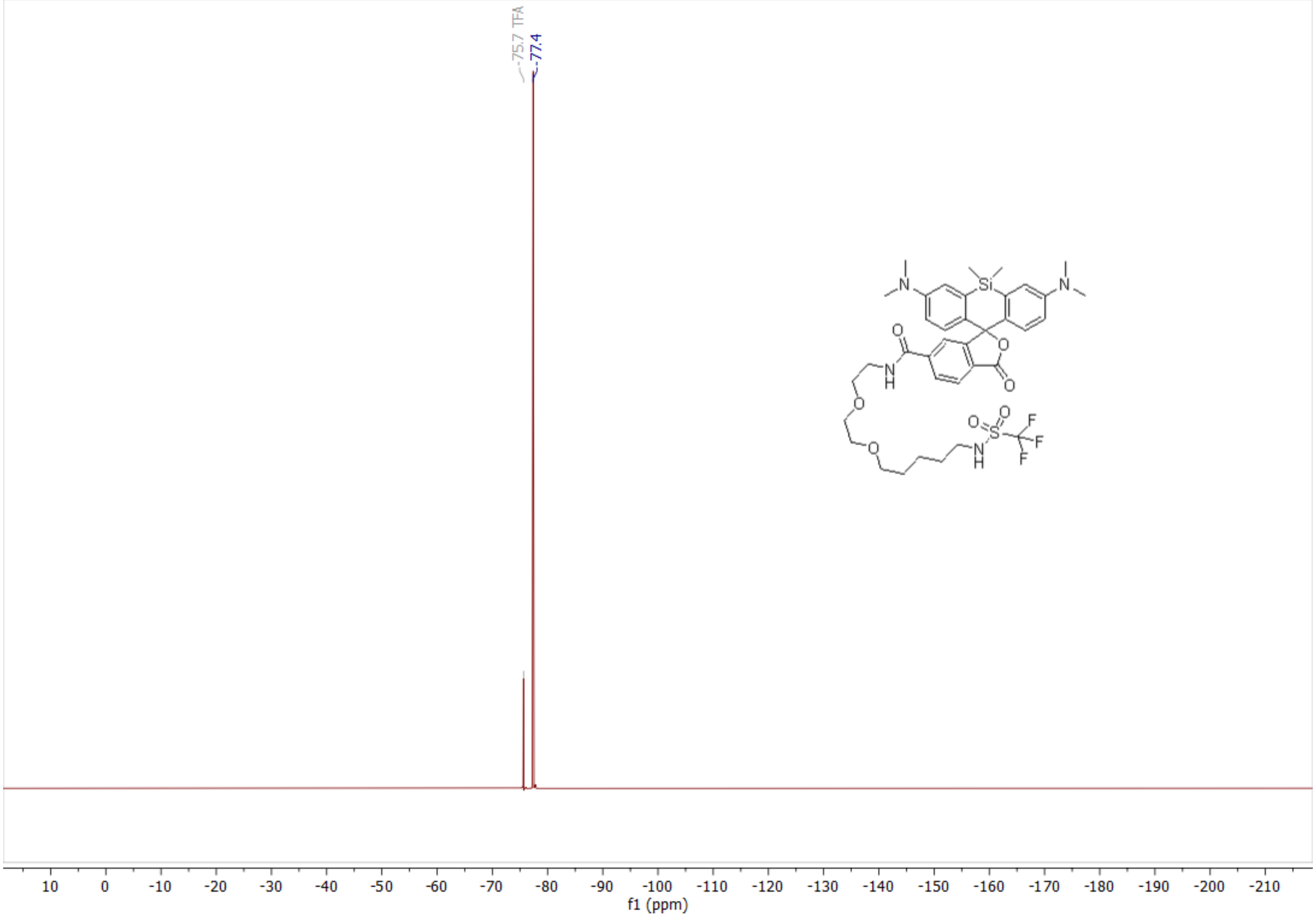
